# Contributions of intra- and extracellular antibiotic degradation to collective β-lactam survival

**DOI:** 10.1101/2024.10.14.618215

**Authors:** Rotem Gross, Muhittin Mungan, Suman G. Das, Melih Yüksel, Berenike Maier, Tobias Bollenbach, J. Arjan G.M. de Visser, Joachim Krug

## Abstract

Collective antibiotic resistance occurs when populations of bacteria survive antibiotic treatments that are lethal to individual bacteria, which affects the efficacy of drug therapies. An important mechanism of collective resistance against widely used *β*-lactams is the production of drug-degrading *β*-lactamases. Here, we integrate experiments with mathematical modeling to understand the collective survival of *Escherichia coli* challenged with cefotaxime. At near-lethal cefotaxime concentrations, we observe complex dynamics, involving initial biomass growth due to filamentation, followed by death, and subsequently growth recovery. We show that production of AmpC, a chromosomal *β*-lactamase, is responsible for cefotaxime degradation, allowing the resumption of cell division in surviving filaments. The detoxification of the environment proceeds through CTX hydrolysis by AmpC in the periplasm of intact cells, as well as extracellularly after cell lysis. Our model predicts the recovery time from molecular parameters, and quantifies the relative roles of periplasmic and extracellular degradation for two strains of *E. coli* that differ in the degree of privatization of AmpC hydrolysis. Our findings suggest that *β*-lactam survival of bacterial infections depends on a combination of intra- and extracellular *β*-lactamase activity, which will likely vary among isolates.

## Introduction

The concept of antibiotic resistance extends far beyond the modern clinical use of antibiotics [1]. Resistance genes have been identified in pristine settings, including the Alaskan soil and ocean floor sediments [2, 3], and *β*-lactamases, key enzymes responsible for resistance against *β*-lactam antibiotics, are thought to have originated billions of years ago [4, 5]. While the roles of antimicrobials and the associated resistance mechanisms in natural ecosystems are unclear, their wide distribution and ancient origins indicate that they may have evolved as a defense against resource competitors and predators [6].

In the clinical context, *β*-lactam antibiotics are the most widely prescribed class of antimicrobial compounds used to treat a variety of infectious diseases [7]. Correspondingly, *β*-lactam resistance through degradation by *β*-lactamases has been a major medical concern for decades [4, 8, 9, 10, 11]. The enzymatic breakdown occurs in the periplasm of gram-negative bacteria, but it can also take place extracellularly if the enzyme is excreted or released from lysing cells [8, 12]. Both modes of degradation reduce the extracellular concentration of the drug, which makes *β*-lactamase production a paradigm of a public good that benefits producing as well as non-producing cells [13, 14, 15, 16, 17, 18, 19, 20], and causes *β*-lactam resistance to be a collective property of the bacterial population [21, 22].

Previous work on the social effects of *β*-lactamase production has mostly considered communities of bacterial strains [12, 13, 17, 23, 24], focusing on the interaction between producers (“helpers”) and non-producers (“beneficiaries”) [14, 16]. Studies of collective *β*-lactamase-induced resistance of isogenic populations have been limited to global inferences about the establishment of colonies from single cells [25, 26, 27] or the inoculum effect, where the minimal inhibitory drug concentration (MIC) increases with cell density [15, 20, 21, 25, 28]. Another phenomenon that has been reported in experimental studies with isogenic populations dating back at least 50 years, is complex growth dynamics at near-lethal *β*-lactam concentrations, where a transient increase in biomass is followed by a decline and a subsequent recovery [8, 23, 29, 30, 31, 32, 33, 34]. However, a mechanistic understanding has only recently begun to emerge.

Here, we combine experiments and modeling to analyse the contributions of periplasmic and extracellular (after the release from lysed cells) *β*-lactamase-mediated antibiotic degradation to the survival of isogenic populations of *Escherichia coli* in liquid culture in the presence of cefotaxime, a third-generation cephalosporin. We will experimentally demonstrate, that the complex growth dynamics arise from an interplay of transient tolerance due to the formation of filamentous cells and breakdown of the antibiotic by AmpC, a chromosomally encoded *β*-lactamase [35] known to contribute to *β*-lactam resistance [36]. Filamentation is a common morphological response to *β*lactam antibiotics [37, 38], which reflects the mode of action of these drugs. *β*-lactams interfere with the synthesis of bacterial cell walls through binding to specific proteins called penicillin-binding proteins (PBPs). Cefotaxime interferes primarily with PBP3, which plays a central role in the divisome, the protein complex involved in cell division. Inhibition of PBP3 obstructs cell division and induces the formation of filamentous bacterial cells [39, 40, 23, 24].

Filamentation delays cell lysis, which leads to a transient increase of biomass by the elongation of filaments [41, 42]. At the onset of lysis of filamentous cells, the population initially collapses, but over a broad range of drug concentrations, the decline is followed by a subsequent phase of recovery. Previous work in which similar dynamics were observed has attributed the recovery to the breakdown of the antibiotic by extracellular *β*-lactamase released by lysing cells [31]. However, enzymatic degradation also occurs intracellularly in the periplasm of intact cells, and the relative contributions of the two modes of breakdown are not known. Disentangling the effects of intra- and extracellular enzymatic activity is important for predicting the impact of *β*-lactamase inhibitors against *β*-lactamase producing bacteria, since these compounds are typically not able to penetrate the periplasm and therefore affect only the extracellular breakdown [43].

In this work, we develop a detailed kinetic model of the intra- and extracellular enzymatic breakdown of cefotaxime, which we couple to the production and decay of biomass obtained from experimental growth curves. We validate the model by comparing predicted and observed recovery times, and use it to estimate the relative contributions of intra- and extracellular breakdown over the time course of the experiment. Experiments are conducted with two common laboratory strains of *E. coli* that differ substantially in the contribution of intra- and extracellular *β*-lactam hydrolysis to the recovery kinetics. A theoretical analysis identifies the conditions under which intra- and extracellular hydrolysis are expected to occur, and points to a generic tradeoff between the private benefit of intracellular *β*-lactamase activity and the contribution to collective survival by the release of the enzyme through lysis.

## Results

### Growth of *E. coli* in the presence of cefotaxime (CTX) shows complex dynamics associated with filamentation, death, and recovery

Growth curves of *E. coli* strain REL606 in the presence of increasing CTX concentrations show qualitative differences (Figure 1A). For low CTX concentrations, biomass increases monotonically, saturating at large times. For concentrations between 0.042 *µ*g/ml and 0.262 *µ*g/ml, the curves exhibit non-monotonic behavior: the initial growth phase is followed by a decay of biomass until around 15h, before a recovery phase with the resumption of growth sets in. Finally, at even higher concentrations the initial growth is followed by a monotonic decay of biomass, and the populations do not consistently recover. Figure 1B summarizes the growth, decay and recovery rates in these three phases as a function of initial CTX concentration (see the SI for details).

**Figure 1:**
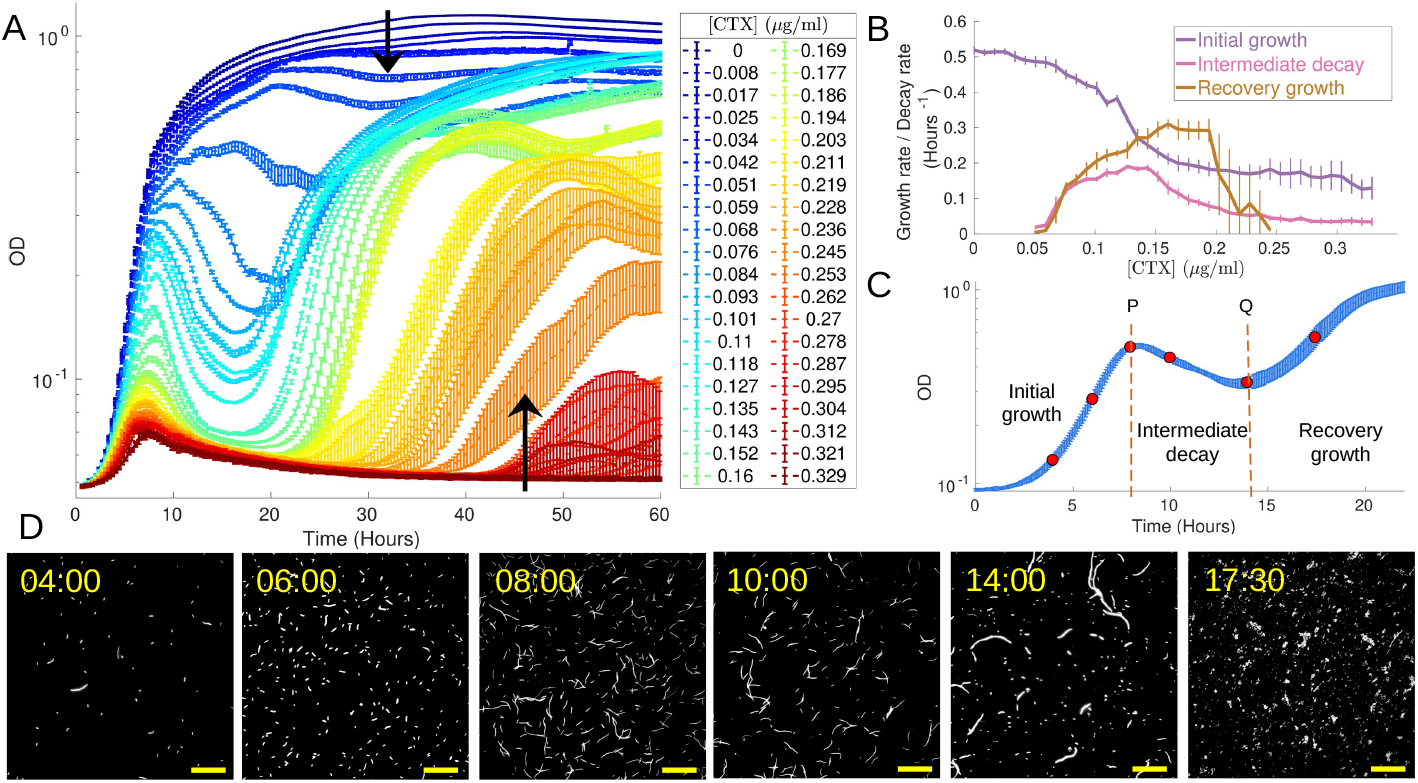
Non-monotonic growth dynamics of *E. coli* REL606 in the presence of cefotaxime (CTX). **A**: Growth curves (measured as optical density, OD, at 600nm) in the presence of varying concentrations of CTX. Each curve shows the average and standard deviation (SD) of OD of 24 replicate cultures initiated with 5 · 10^5^ cells/ml. Non-monotonic growth, involving initial growth, decay and recovery, is observed at CTX concentrations between 0.042-0.262 *µ*g/ml (range boundaries are indicated by black arrows). Beyond a CTX concentration of 0.262 *µ*g/ml, large fluctuations between replicate cultures occur and are accompanied by recovery times exceeding 40h. **B**: Estimates of the average and SD of the maximum growth, decay and recovery rate, attained in the three regimes of the OD growth curves shown in panel A, as a function of CTX concentration (see SI and Fig. S6 for details on the feature extraction). **C**: Mean and SD growth curves of 16 replicate cultures grown at 0.08 *µ*g CTX/ml. The vertical dashed lines mark the initial OD peak *P* and minimum *Q*, separating the three growth phases – growth, decay, and recovery. **D**: Microscope images sampled from the time points marked in red in panel C, showing increasing numbers of filamentous cells during initial growth and dominance of normal-size cells during recovery. The yellow length bar corresponds to 100 *µ*m.

To elucidate the origins of these complex growth dynamics, we first analyzed samples from different time points of populations growing at 0.08 *µ*g CTX/ml under the microscope (Figs. 1C and D). Until biomass peak *P*, cells become increasingly elongated into filaments. Between time points *P* and *Q*, when the biomass is decaying, the number of elongated cells decreases in consecutive images. Around *t* = 14h (point *Q* in panel C), the recovery phase sets in. Already at this time, normal-size cells are visible, which by *t* = 17.5h have taken over the whole population.

The close similarity of the replicate growth curves measured at the same CTX concentration in Figs. 1A and C makes it unlikely that spontaneous mutations cause the recovery of growth at intermediate CTX concentrations. Regrowth experiments with 80 replicate cultures essentially rule out this possibility (see Fig. S1), which was confirmed by whole genome sequencing (see Materials and Methods).

### Growth recovery can be explained by filaments resuming cell division when CTX is removed

We hypothesize that the resumption of cell division during recovery is due to the reduction of CTX in the medium. To see whether normal size cells survive CTX stress or filamentous cells resume cell division, we observed cell growth in a microfluidic chamber. We first treated the bacterial population with a standard LB medium containing 0.2 *µ*g CTX/ml (causing non-monotonic growth) for 2.5 h, followed by the same growth medium without CTX for 20 h (Fig. 2A). By observing one or a few cells per field of view over time, we were able to follow their response to the removal of CTX. As expected, we found that during the initial flow with CTX, the cells formed filaments (Fig. 2B). About 50 minutes after the end of the treatment with CTX, we observed that filamented cells started to divide again (see Fig. 2C and Supplemental Movie). We, therefore, conclude that the transition from growth decay to growth recovery in Fig. 1A is due to the onset of division of filamented cells. In agreement with Ref. [44], we observe that the resumption of cell division of the filaments can occur along any septum and there is no preference for budding off normal-sized cells from the tips of filaments.

**Figure 2:**
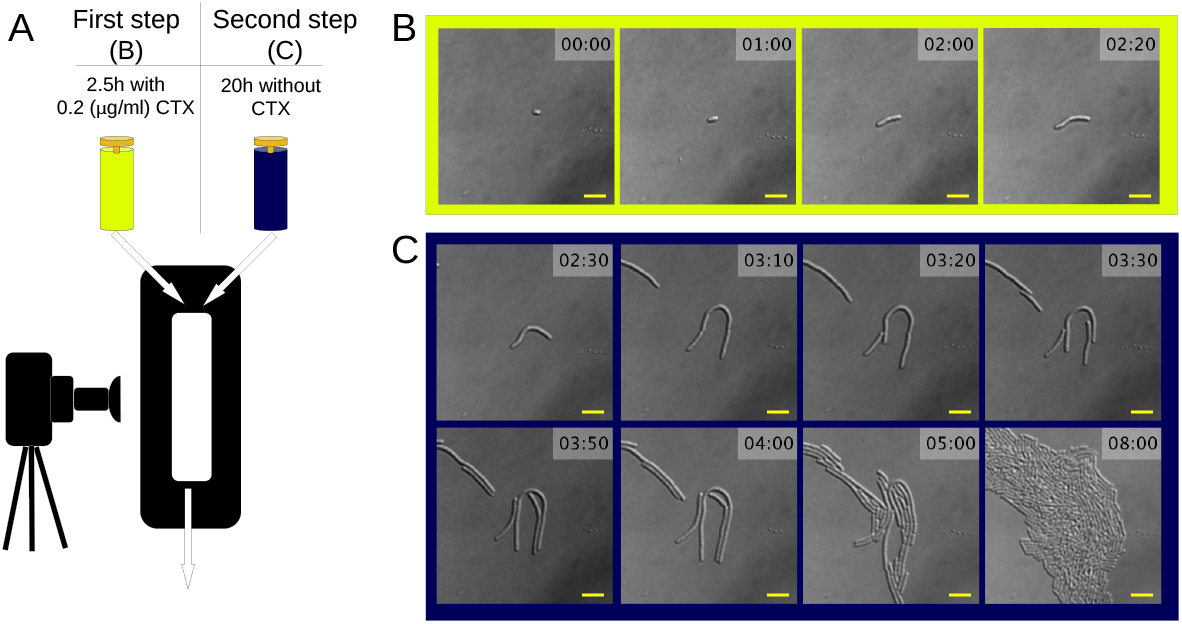
Filamentous cells resume cell division when CTX is removed. **A**: *E. coli* REL606 was grown in a microfluidic flow chamber connected consecutively to two syringes, enabling the supply of nutrient medium with and without CTX. During the first 2.5h, nutrient medium with 0.2 *µ*g CTX/ml (lemon green) was injected, followed by 20h of CTX-free nutrient medium (dark blue). **B, C**: Microscope images from the two phases of the experiment (B: medium with CTX; C: medium without CTX). During exposure to CTX, the cell in the center of view starts to filament. Filamentation continues until 3h10, i.e. 40 min after exposure to CTX was stopped. Cell division accompanied by elongation of filaments occurs between *t* = 3h20 and *t* = 4h, resulting in dividing cells that are still larger than their normal size. However, by *t* = 8h all cells are normal-sized. The yellow length bar corresponds to 10 *µ*m.

### Spent medium from CTX-treated cultures accelerates growth resumption

The microfluidic chamber experiment suggests that growth recovery in the batch culture experiments (Fig. 1) is due to the removal of CTX by the growing cells. To determine the remaining amount of CTX at the onset of the recovery phase in the batch cultures, we designed three spent medium experiments (see Materials and Methods). In each experiment, we grew *E. coli* REL606 for 14 h in LB medium at a reference concentration of 0.08 *µ*g CTX/ml, which corresponds to the lower range of concentrations for which we see filamentation and complex growth (see Fig. 1C). We then filtered out the cells and mixed the resulting spent medium with freshly made LB medium consisting of nutrients and a prescribed amount of CTX. The amount of CTX in the added fresh medium was adjusted such that, under the assumption that all CTX has been removed in the spent medium fraction, the CTX concentration in the mixture reaches a prescribed nominal value [CTX]_*m*_. If not all CTX had been removed from the spent medium fraction by the bacteria, the actual concentration in the mixed medium would thus exceed [CTX]_*m*_.

In the first experiment, growth in the absence and presence of 45% of spent medium was compared for four [CTX]_*m*_ (Fig. 3A). Whereas in the absence of CTX, spent medium does not affect growth as expected, the spent medium unexpectedly *enhanced* growth in the presence of CTX, indicating protection beyond the complete removal of CTX in the spent medium fraction. The second experiment was a repetition of the first experiment, but with more variation in the spent medium fraction *ϕ*, nominal concentrations [CTX]_*m*_, and using sonication during the preparation of the spent medium to maximize the release of protective agent from the cells. The area under the growth curves (AUC) shows a negative impact of spent medium in the absence of CTX, presumably due to nutrient depletion, but for CTX concentrations above the concentration used to produce the spent medium, growth increases systematically with spent medium fraction (Fig. 3B). We also demonstrated the protectiveness of the spent medium on agar medium (Fig. S2). In the third experiment, we asked whether the protective effect of spent medium is sensitive to heat, which would suggest it is conferred by a protein. To test this, the spent medium was subjected to a heat treatment prior to the mixing with fresh medium. The growth curves show that heating the spent medium reduces its protection ability in the presence of CTX (Fig. 3C), but has no effect in fresh medium (Fig. 3D), suggesting that a protein secreted by the bacteria is the cause of protection.

**Figure 3:**
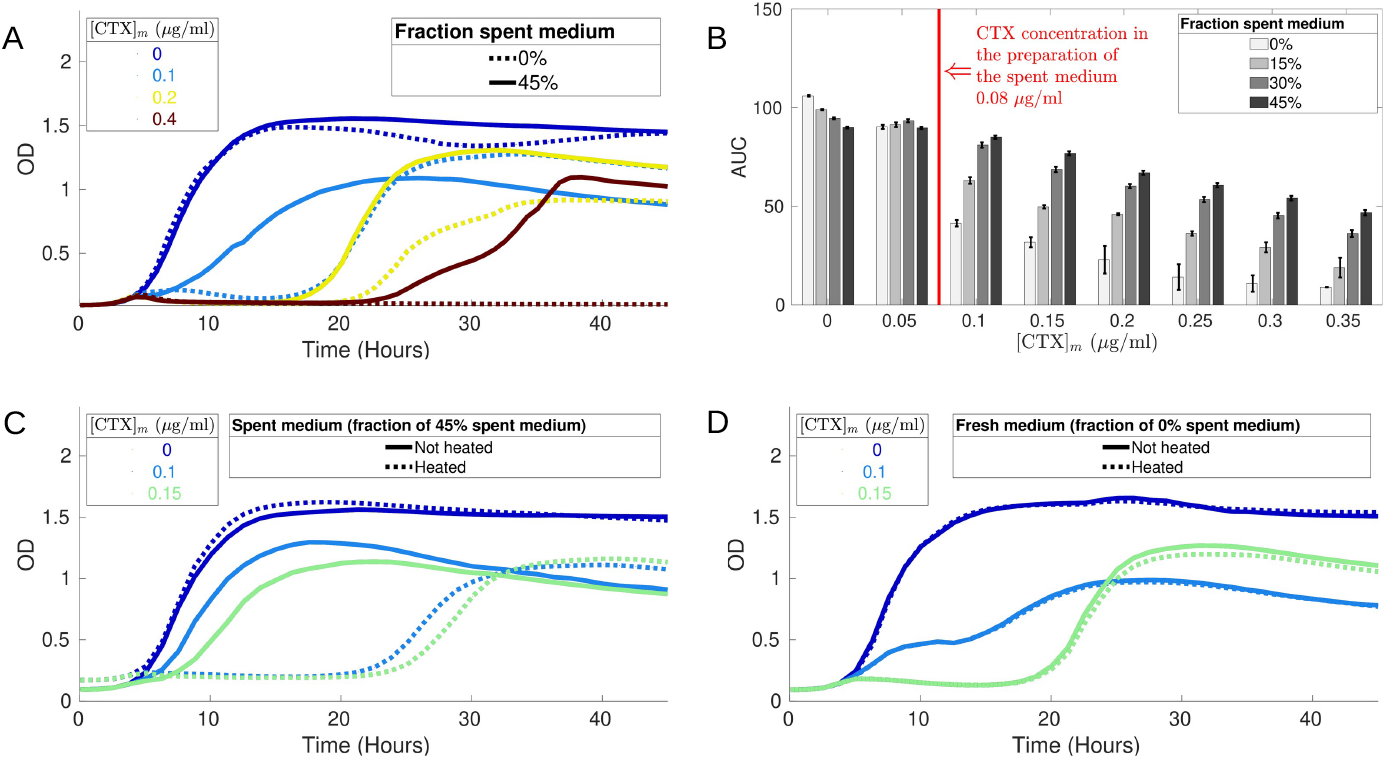
Spent medium from CTX-treated *E. coli* REL606 cultures is protective. **A**: Growth curves at four [CTX]_*m*_ of cultures containing 0% (dashed lines) or 45% (solid lines) spent medium. The spent medium was extracted from overnight cultures grown in the presence of 0.08 *µg* CTX/ml. [CTX]_*m*_ are nominal initial concentrations in mixed cultures under the assumption that all CTX in the spent medium fraction was removed. **B**: Bacterial growth (measured by the area under the growth curve, AUC, during 30h) at different spent medium fractions *ϕ* (harvested after 10 s of sonication to lyse the cells) and [CTX]_*m*_. **C**: The protective effect of 45% spent medium in the presence of CTX disappears after heating (98^°^C for 30 minutes). **D**: Heating has no effect on growth in fresh medium.

### AmpC *β*-lactamase expression accelerates growth recovery in the presence of CTX

Based on the results presented in the preceding section, it appears that the protective effect of the spent medium is related to a protein released by the cells. A likely candidate is the *β*-lactamase AmpC, since REL606 does not contain plasmids and AmpC is the only chromosomally-encoded enzyme with *β*-lactamase activity (https://ecoliwiki.org/colipedia/index.php/Category:Gene_List:REL606).

To test this hypothesis, we performed a parallel set of experiments using the *E. coli* K12 strain, for two reasons. First, the availability of the ASKA [45] and KEIO [46] strain collections for K12 (see Materials and Methods for details) allowed us to vary the expression level of *ampC*, which K12 expresses at a basal level like REL606 [47, 35]. Second, chemical rate constants and other parameters affecting the hydrolysis of CTX by K12’s AmpC have been determined [48, 49], which implies that we can link the delay times marking the onset of the recovery phase to the reaction kinetics of CTX hydrolysation by quantitative modeling.

Growth behavior of *E. coli* K12 Δ*lacA* (Fig. S3) is qualitatively similar to REL606 (Fig. 1A), except that non-monotonic growth sets in at a fivefold lower CTX concentration and the transient decay of biomass is shallower, suggesting lower lysis rates. In order to assess the effect of AmpC expression on growth recovery, we used strain JW4111 from the ASKA collection. This strain contains *ampC* under an inducible promoter on a multicopy plasmid (in addition to its copy in the chromosome), allowing regulated overexpression by varying the concentration of inducer IPTG in the medium. We grew JW4111 with three different concentrations of IPTG for 24 h without CTX, filtered out the cells, mixed the remaining spent medium with fresh medium and then let the bacteria grow in the mixed medium in a CTX concentration gradient in the absence of IPTG. Expression of AmpC in the spent medium strongly accelerates growth recovery and causes complete protection for the highest expression level at CTX concentrations far beyond concentrations that are lethal in the absence of spent medium (Fig. 4A).

**Figure 4:**
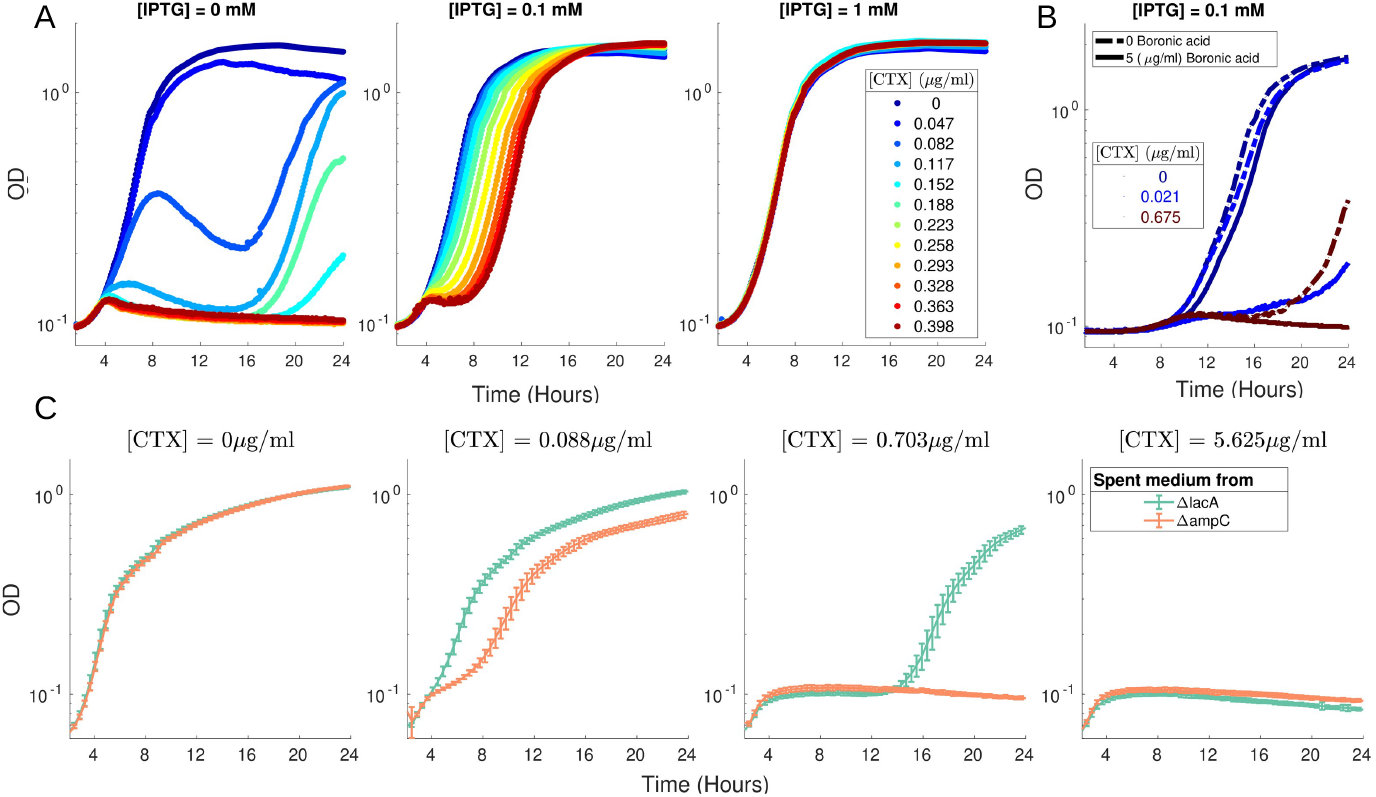
AmpC expression accelerates growth recovery in CTX. **A**: Increased expression of AmpC from a plasmid accelerates growth recovery and allows for monotonic growth at CTX concentrations that fully inhibit growth in the absence of AmpC induction. The three panels show growth curves of *E. coli* JW41111 (ASKA collection, see Materials and methods) in spent media harvested from overnight cultures, grown in the absence of CTX, of the same strain without and with two levels of expression of AmpC induced by IPTG. Note that the growth curves in the leftmost panel display higher OD and a more pronounced non-monotonicity than the corresponding curves for K12 Δ*lacA* in Fig. S3. This likely reflects higher AmpC expression due to its higher copy number and leakiness of the inducible promoter. **B**: The addition of boronic acid, which binds to AmpC and inhibits its function, removes the protective effect of AmpC at an intermediate expression level in the presence of CTX. **C**: Growth of K12 Δ*lacA* in fresh medium mixed with 45% spent medium from overnight cultures of strains K12 Δ*lacA* or Δ*ampC* in the absence of CTX, at four different CTX concentrations.

We then repeated the experiment at intermediate AmpC induction by splitting the spent medium into two fractions. Adding the *β*-lactamase inhibitor boronic acid to one fraction led to a substantial reduction of growth (Fig. 4B). To confirm the causal role of AmpC in CTX removal and growth recovery, we performed two further experiments. First, we over-expressed each *amp* gene separately, using strains from the ASKA collection and measured the growth on a CTX gradient. The data shown in Fig. S4 clearly demonstrate that only over-expression of AmpC confers increased CTX resistance. Second, we harvested the spent medium from sonicated overnight cultures of the KEIO deletion strains Δ*ampC* and Δ*lacA*, and grew K12 Δ*lacA* in mixed media with varying CTX concentrations. At intermediate CTX concentrations, spent medium from the strain Δ*lacA*, which expresses AmpC, enhances growth relative to that in the spent medium extracted from Δ*ampC* (Fig. 4C). Taken together, these findings firmly validate our hypothesis that the observed protection is due to CTX breakdown by AmpC *β*-lactamase.

### A kinetic model of CTX hydrolysis predicts the recovery time

The scenario emerging from our results is that the observed growth recovery following decay is due to hydrolysis of CTX by AmpC. We hypothesize that growth resumes once the drug concentration has been reduced below a threshold level. To corroborate this scenario, we developed a model of CTX hydrolysis by AmpC in order to predict recovery times and compared these to observations.

We assume that CTX is hydrolysed by AmpC at two locations: in the periplasm after diffusion through porins in the outer membrane, and in the extracellular medium after the release of AmpC by cell lysis. We use the experimentally obtained growth curves to estimate the amount of non-lysed and lysed biomass at any given time during the experiment, which we then take as input to calculate (i) the amount of AmpC production (by the non-lysed biomass), (ii) the fraction released into the medium upon lysis, and coupled to this, (iii) the total amount of CTX hydrolysed over time by taking into account its reaction kinetics. Similar modeling approaches have been used by other groups [15, 28, 42, 34]. Here, we aim for a maximally parsimonious description that explains the observed kinetics using a minimal set of assumptions and parameters. A detailed discussion of the model and its predictions is provided in the SI. In the following, we outline the main ingredients and present our results.

We consider first the hydrolysation of CTX in the periplasm [48, 49, 50]. An equilibrium concentration inside the periplasm is achieved when the diffusion of CTX molecules into the periplasm is balanced by their hydrolysis by AmpC. The CTX concentration difference across the outer membrane is then given by the Zimmermann-Rosselet equation [51],

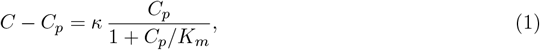

which relates the extracellular and periplasmic CTX concentrations, *C* and *C*_*p*_, via the dimension-less parameter

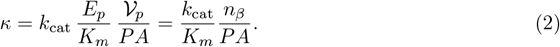

Here, *K*_*m*_ and *k*_cat_ are the Michaelis-Menten reaction parameters describing CTX hydrolysis, *P* is the permeability constant of the outer membrane, *A* is the surface area of *E. coli*, 𝒱_*p*_ is the periplasmic volume, *E*_*p*_ is the enzyme concentration inside the periplasm and *n*_*β*_ = *E*_*p*_ 𝒱_*p*_ is the number of AmpC molecules per normal-size cell. The dimensionless parameter *κ* is the ratio of the rates of hydrolysation and influx of CTX in the periplasm.

Equation (1) shows that when *κ* is small, the CTX concentrations outside the cell and within the periplasm are nearly equal. Conversely, when *κ* is large, the concentration *C*_*p*_ of CTX inside the periplasm is much lower than outside. In this sense, *κ* quantifies the degree of privatization of *β*-lactam hydrolysis. Importantly, the privatization of the hydrolysis should not be confused with the privatization of the enzyme. For intact cells, the enzyme is localized in the periplasm and remains fully private. In contrast, its function of *β*-lactam degradation is ‘leaky’, in the sense that it reduces environmental concentrations and hence benefits both producing and non-producing cells, but to different degrees [16, 19]. In the absence of other resistance mechanisms, such as efflux pumps, the periplasmic concentration of non-producing cells would be equal to the external concentration *C > C*_*p*_, and they are therefore more strongly affected by the antibiotic.

We estimate the time-dependence of the CTX concentration in the extracellular medium by taking into account the reaction kinetics as well as the bacterial growth kinetics, which we infer from the experimental OD curves. Our working assumption is that the OD signal is proportional to the total live biomass, including filaments as well as normal-size cells (see [34] for a detailed discussion of the relation between OD and colony counting assays). We are concerned with the regime in which the change of biomass is non-monotonic (Fig. 1). Initially the biomass increases at an average growth rate *g* up to some time *t*_*P*_ at which the OD reaches its maximum. For *t > t*_*P*_, we assume that lysis sets in with a fixed rate *γ*, such that the net biomass decay rate is given by *γ* − *g*′, where *g*′ is the rate at which cells continue to grow beyond *P*. For simplicity, we assume that *g* = *g*′, which implies that the biomass growth rate remains constant. Figure S7A illustrates the extraction of the average initial growth and decay rate from empirical OD curves. The resulting estimates of the rates *g, γ* − *g* and *γ* are shown as a function of the initial CTX concentration in Fig. S8A and B. Incorporating periplasmic CTX degradation by live cells as well as extracellular degradation by AmpC released after cell lysis, the time evolution of the total amount of CTX in the medium is then given by the differential equation (S18), which is solved numerically to predict the recovery time.

For *E. coli* K12, Nikaido and collaborators have measured the permeability *P*, as well as the reaction constants for CTX hydrolysation by AmpC [48, 49]. We therefore consider the behavior of this strain first. Similar to REL606, the non-monotonic growth of *E. coli* K12 exhibits three phases (Fig. S3). We assume that the onset of the recovery phase is determined by the time at which the extracellular CTX concentration falls below the value for which the OD growth is still monotonic, i.e. *C* = 0.015 *µ*g/ml for *E. coli* K12. All parameters in our model can be inferred from experiments, except for the number *n*_*β*_ of AmpC molecules per normal-size cell. We therefore take *n*_*β*_ as the single adjustable model parameter to fit the data. Figure 5A shows a comparison of the empirically observed recovery times *τ*_rec_ = *t*_*Q*_ − *t*_*P*_ for *E. coli* K12 with the predicted recovery times from our model. For *n*_*β*_ = 700 AmpC molecules per cell (in line with previous estimates [52, 53]), we obtain recovery times that follow the data points reasonably well over a broad range of CTX concentrations. In view of the minimal character of our model, which does not include physiological changes in the production rate of AmpC or the cell wall permeability, the agreement is rather satisfactory.

**Figure 5:**
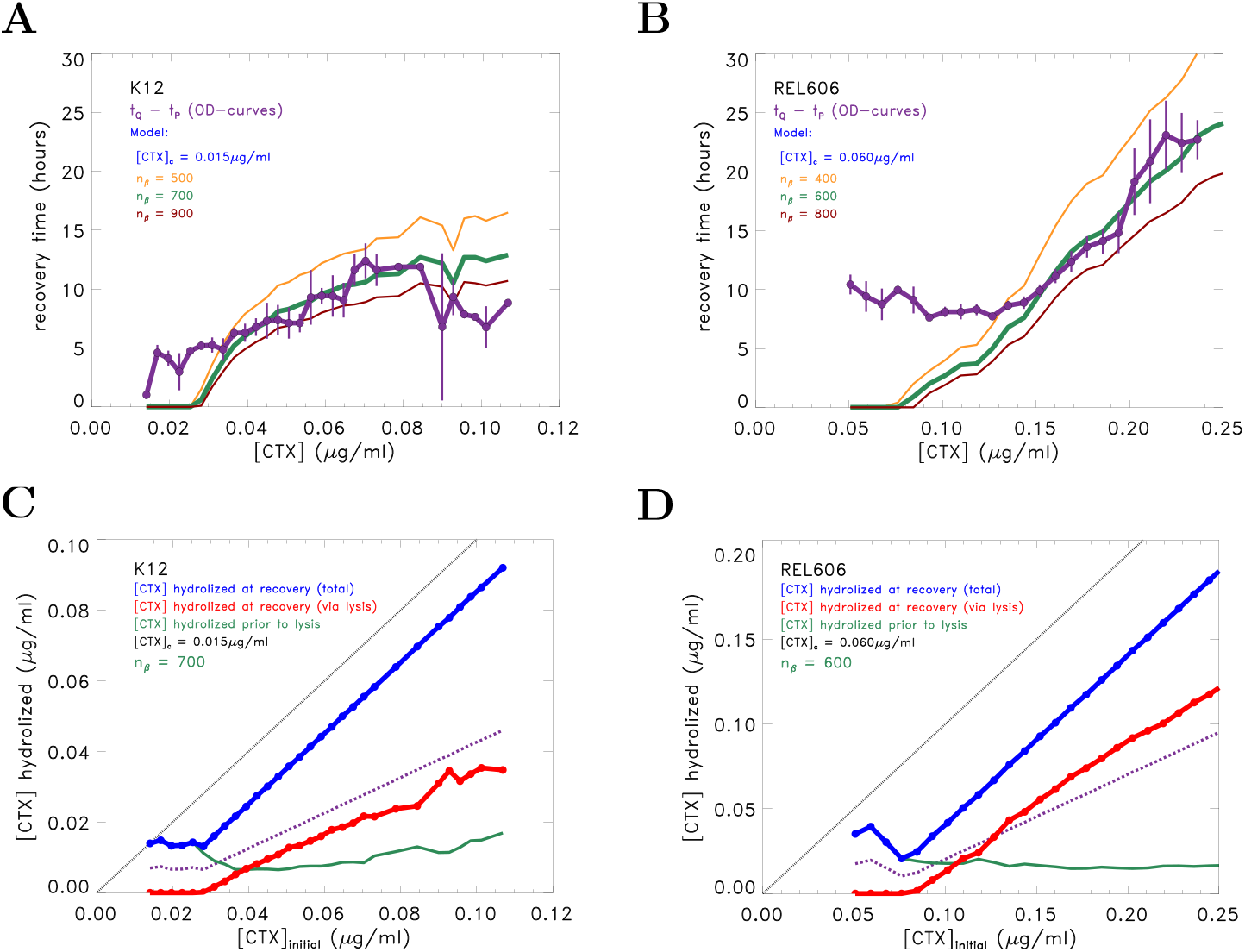
Model predictions of the time required for AmpC to hydrolyse CTX are consistent with observed recovery times. **A**: Comparison of recovery times of *E. coli* K12 inferred from the growth curves (black lines and data points) with predictions from the kinetic model, using the number *n*_*β*_ of AmpC molecules per normal-size cells as a free parameter. The best agreement is obtained for *n*_*β*_ = 700. **B**: The same comparison for *E. coli* REL606, showing reasonable agreement for *n*_*β*_ = 600. **C**: Amount of CTX hydrolysed until recovery as a function of initial drug concentration for *E. coli* K12. The diagonal dashed line is a guide to the eye representing the case that all CTX has been hydrolysed at recovery. The assumed recovery condition is [CTX]_hydrolized_ = [CTX]_initial_ − [CTX]*, with [CTX]* = 0.015*µ*g*/*ml, as illustrated by the solid blue line for [CTX]_initial_ ≥ 0.030*µ*g*/*ml. The green curve shows the amount of CTX hydrolysed during the initial growth phase prior to the onset of lysis. The red curve is the contribution due to hydrolysis in the extracellular medium as computed from the model and the experimental OD-curves. The purple curve depicts the case when the periplasmic and extracellular contributions to hydrolysis are equal. For *E. coli* K12, more than half of the CTX is hydrolysed within the periplasm. **D**: Amount of CTX hydrolysed until recovery for *E. coli* REL606. Beyond [CTX]_initial_ = 0.13*µ*g*/*ml, extracellular hydrolysis contributes more than half of the total hydrolysed CTX.

From our estimate of *n*_*β*_ = 700, we find using (2) that *κ*_*K*12_ = 0.39 (see Materials and Methods). To determine the CTX concentration gradient across the outer membrane from Eq. (1), we use the cross-over from monotonic to non-monotonic growth as an indicator that CTX concentrations within the periplasm have reached levels at which cell growth starts to be affected. As mentioned above, this happens around concentrations of *C* = 0.015 *µ*g CTX/ml. With the estimated value of *κ*_*K*12_, we find from (1) that the corresponding concentration inside the periplasm is *C*_*p*_ = 0.011 *µ*g/ml, which is comparable to the CTX concentration outside. We conclude that for *E. coli* K12 at sub-lethal drug concentrations, the transport of the drug across the outer membrane is sufficiently efficient to maintain similar CTX concentrations inside and outside the cell.

We next apply the same approach to *E. coli* REL606. For this strain, non-monotonic OD growth sets in at CTX concentrations above *C* = 0.060 *µ*g/ml (Fig. 1). Assuming that this occurs at the same periplasmic concentration of *C*_*p*_ = 0.011 *µ*g/ml as for *E. coli* K12, the corresponding value is *κ*_REL606_ = 5.0, which is about 13 times larger than *κ*_K12_. From Eq. (2) we see that this could be a result of a difference in enzyme activity, the number of enzyme molecules *n*_*β*_ or permeability of the outer membrane. We were not able to find published estimates of *P*, *k*_cat_ and *K*_*m*_ for REL606. However, since the value of *κ* is fixed, we have again only one adjustable parameter, which we choose to be *n*_*β*_.

With these assumptions made, the recovery times for REL606 can be predicted from the OD curves in the same way as for K12, as shown in Fig. 5B. The best agreement with the empirical data occurs when *n*_*β*_ = 600. The fits of the recovery times thus yield a comparable number of enzyme molecules for both strains. This suggests that the larger *κ*-value in REL606 reflects a lower outer membrane permeability or a higher enzyme efficiency, or a combination of both effects.

### *E. coli* strains REL606 and K12 differ in the degree of periplasmic versus extracellular CTX hydrolysis

As we have seen, the magnitude of the CTX concentration gradient across the outer membrane is governed by *κ*, which is about 13 times larger for *E. coli* REL606 than for K12. We next use our modeling approach to examine how this difference affects the relative contributions of periplasmic and extracellular enzymatic activity to CTX removal.

Recall that the hydrolysis rate is proportional to the product of enzyme and substrate concentrations (see Eq. S2). Thus under conditions where AmpC concentrations are higher inside cells and CTX concentrations are higher outside cells, the release of enzymes into the medium will generally speed up hydrolysis. For *E. coli* REL606, we therefore expect lysis to contribute more strongly to the overall breakdown compared to *E. coli* K12. Our model permits us to determine the amounts of CTX hydrolysed in the periplasm and in the medium by separately integrating the corresponding terms on the right hand side of Eq. (S18). Figure 5C,D shows the result of this analysis. For *E. coli* K12, we see that for the full range of initial CTX concentrations most of the CTX is hydrolysed in the periplasm, while for *E. coli* REL606, this is only the case for low initial concentrations. For initial CTX concentrations above 0.13 *µ*g/ml, increasingly larger amounts of CTX are hydrolysed in the medium.

Figures S8C and D illustrate how the enzymatic breakdown in the two strains progresses over time. The initial concentrations chosen, along with their empirical growth, decay and lysis rates, are indicated by the circled data values in Fig. S8A and B. For *E. coli* K12, we see that at all times, even beyond the onset of recovery, the breakdown of extracellular CTX due to hydrolysis in the periplasm exceeds the contribution due to lysis. This is in contrast to the behavior of *E. coli* REL606. Here, the contribution of lysis starts to dominate well before the onset of recovery.

### Factors determining the contributions of lysis and privatization to collective recovery

The analysis presented above suggests that the K12 and REL606 strains differ in two respects. On the one hand, *β*-lactam degradation by REL606 is more private than by K12 [*κ*_REL606_ ≫ *κ*_K12_]; on the other hand, REL606 displays more cell lysis in the regime of biomass decay [(*γ*−*g*)_REL606_ ≫ (*γ*− *g*)_K12_, see Fig. S8A,B. To gain a general understanding of how these factors contribute to collective survival under antibiotic stress, we now turn to an in-depth investigation of the mathematical model.

For convenience, we introduce dimensionless concentrations *a* = *C/K*_*m*_ and *a*_*p*_ = *C*_*p*_*/K*_*m*_, and assume that both *a* and *a*_*p*_ are much smaller than 1. The conditions under which this is true are specified in the SI, where we also indicate how the analysis can be extended to other regimes. The evolution equation for the dimensionless extracellular *β*-lactam concentration then becomes

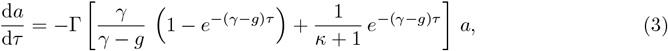

where *τ* = *t* − *t*_*P*_ is the time after the onset of biomass decay at time *t*_*p*_ and

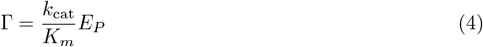

sets the overall scale of the rate of *β*-lactam breakdown. Here *E*_*P*_ is the concentration of *β*-lactamase in the medium that would have been attained if all the cells present at time *t*_*P*_ had lysed at once. It is proportional to the biomass concentration at time *t*_*P*_ as well as to the number of enzyme molecules per cell. The first and second term inside the rectangular brackets of (3) describes the contributions due to hydrolysis in the extracellular medium and the periplasm, respectively. The extracellular contribution is amplified by a factor 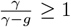 reflecting the turnover of biomass generation and lysis, and the periplasmic contribution is reduced by a factor 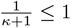 that quantifies the partial privatization of the hydrolysis. The ratio of the two factors defines the dimensionless parameter

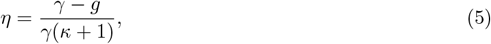

which describes the strain’s propensity for extracellular versus periplasmic CTX hydrolysis. Interestingly, this parameter turns out to be of a similar order (*η* ~ 0.05) for both strains over the range of concentrations we considered.

The main features of the degradation process can be inferred from (3) without detailed analysis. Since the number of intact cells declines over time, periplasmic breakdown dominates at short times [(*γ* −*g*)*τ* ≪ 1] and extracellular breakdown dominates at long times [(*γ* −*g*)*τ* ≫ 1]. The transition between the two regimes occurs at a time *τ*_*c*_ obtained by equating the two terms in the square brackets on the right hand side of Eq. (3), which yields

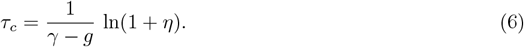

To determine the contribution of extracellular hydrolysis to recovery, this time has to be compared to the time scale *τ*_rec_ at which a sufficient amount of the drug has been hydrolysed to allow for population recovery. The latter depends on the overall breakdown rate (4) as well as on the ratio of the initial drug concentration *a*_0_ to the concentration *a*\* at which recovery sets in. Assuming for simplicity exponential breakdown at rate Γ, we obtain the estimate

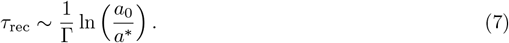

Increasing Γ by, for example, increasing the efficiency of the enzyme or the initial cell density, decreases *τ*_rec_ compared to *τ*_*c*_ and, therefore, reduces the extracellular contribution, whereas an increase of the initial drug concentration *a*_0_ has the opposite effect. On the other hand, increasing the degree of privatization of *β*-lactam hydrolysis decreases *η* according to (5) and reduces *τ*_*c*_ relative to *τ*_rec_, which increases the contribution to recovery from extracellular hydrolysis. Since the parameter *η* in (6) is of a similar order for the two strains, we conclude that the fact that CTX hydrolysis in K12 (REL606) is predominantly periplasmic (extracellular) is primarily due to the difference in the biomass decay rates for the two strains (Fig. S8A,B).

For the explicit solution of (3) it is convenient to introduce the dimensionless parameter

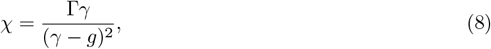

which is a decreasing function of the lysis rate *γ*. In terms of the parameters *η* and *χ*, the solution of (3) can be compactly written as

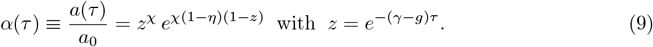

Here, *z* denotes the fraction of the initial biomass that remains at time *τ*. Recovery sets in when *a*(*τ*_rec_) = *a*\* or

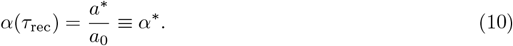

Based on the solution (9), the total amount of *β*-lactamase that has been hydrolysed at recovery can be decomposed into a periplasmic and an extracellular contribution according to 1 − *α*\* = *α*_peri_ +*α*_medium_. The case where equal amounts of CTX are hydrolysed in the extracellular medium and periplasm corresponds to a choice of parameters *α*\*, *χ* and *η* such that 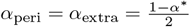. This defines a two dimensional surface in the three dimensional space of parameters.

Figure 6A shows cross-sections of this surface for different choices of *η*. Each curve separates the (*α*\*, *χ*)-plane into regions. In the region above (below) the curve, the contribution of hydrolysis in the periplasm is larger (smaller) than that of hydrolysis in the extracellular medium. We conclude from the figure that a transition from primarily periplasmic to primarily extracellular hydrolysis occurs upon (i) an increase of the initial *β*-lactam concentration (decrease of *α*\*), (ii) a decrease of the rate of *β*-lactam hydrolysis (decrease of *χ* via an decrease of Γ or (iii) an increase of the rate of cell lysis (decrease of *χ* via an increase of *γ*). Moreover, with increasing privatization of hydrolysis (decreasing *η*) the region where extracellular hydrolysis dominates grows.

**Figure 6:**
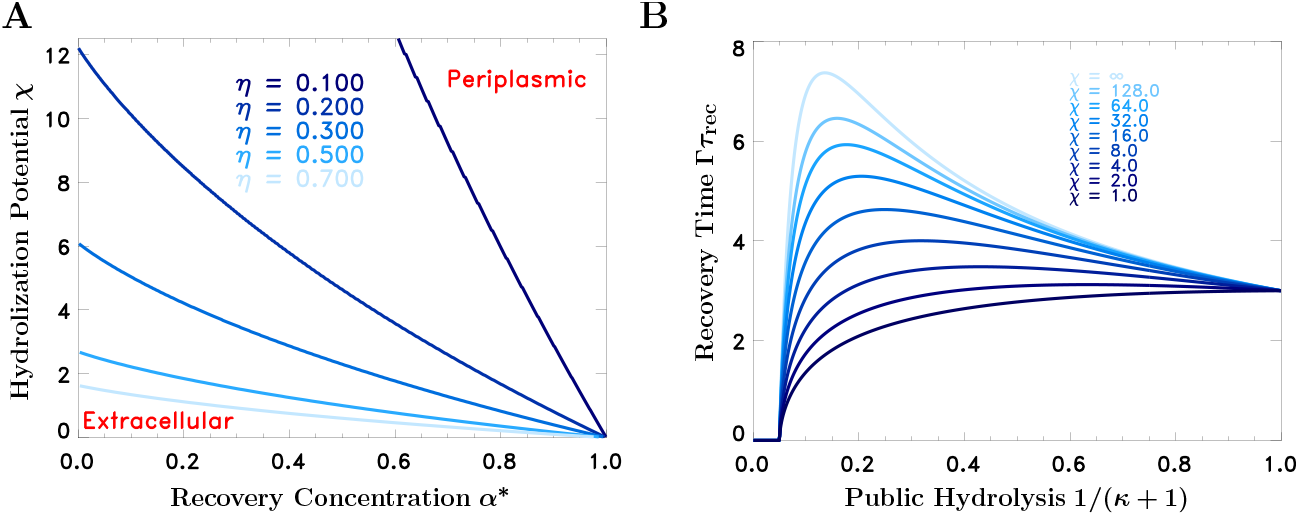
Theoretical analysis of the contributions of lysis and privatization to collective recovery. **A:** The curves separate regions in the plane of dimensionless parameters *α*\* and *χ* for which hydrolysis in the periplasm (above the curves) dominates over hydrolysis in the extracellular medium (below the curves). The parameter *χ* defined in Eq. (8) provides a measure for the hydrolysation potential of the population. The second parameter *α*\* is the ratio of the concentration *a*\* at which recovery sets in to the initial drug concentration *a*_0_, and quantifies the harshness of the environmental challenge. With increasing hydrolysation potential *χ*, the population moves towards the region of predominantly periplasmic breakdown, whereas increasingly harsh environments (decreasing *α*\*) require more extracellular hydrolysis and cell lysis. The curves additionally depend on the degree of privatization *κ* through (5). With increasing privatization (decreasing *η*), extracellular hydrolysis becomes more important. **B:** The panel shows the recovery time *τ*_rec_ as a function of privatization parameter 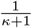 for a scenario where the periplasmic concentration 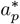 at which recovery sets in is kept constant at 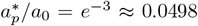. For sufficiently large values of *χ*, the recovery time depends non-monotonically on the degree of privatization. The curve for *χ* → ∞ shows the expression (S35).

### A tradeoff between individual and collective survival

The degree of privatization of *β*-lactam hydrolysis affects the recovery kinetics in two ways. On the one hand, the contribution of periplasmic hydrolysis in Eq. (3) is proportional to 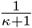. All else being equal, increasing hydrolysis privatization (increasing *κ*), therefore, slows down the degradation process, because it reduces the efficiency of the periplasmic channel. On the other hand, increasing *κ* also reduces the periplasmic drug concentration relative to the external concentration, which implies that recovery will set in at larger external drug concentrations, hence earlier in time.

To explain this last point, we recall that in the analysis of the recovery kinetics of *E. coli* REL606 and K12, we have assumed (following common practice [9]) that recovery begins when the periplasmic concentration falls below a critical value 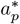, which is set by intrinsic properties of the cells and, therefore, takes the same value for both strains. The corresponding external concentration required for recovery

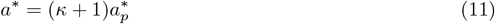

is an increasing function of *κ*. Since the reduction of the periplasmic drug concentration is the primary mechanism through which *β*-lactamase production contributes to the resistance of single cells [15], whereas the recovery time describes the resilience of the population [31], we argue that these two conflicting trends constitute a tradeoff between individual and collective survival.

This tradeoff may lead to a non-monotonic dependence of the recovery time *τ*_rec_ on the degree of privatization of *β*-lactamase mediated hydrolysis. Figure 6B shows the product 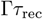 as a function of 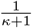, obtained by evaluating (10) using (11) for a fixed value of 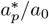 and different values of *χ*. For simplicity it is assumed that 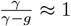, i.e. *γ* ≫ *g*, which implies that *χ* = Γ*/γ*. The relationship between *τ*_rec_ and 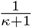 is non-monotonic for sufficiently large *χ*. In this regime an increase of the degree of privatization *κ* increases the time to recovery, and thus decreases collective resilience, in favor of increased single-cell resistance. On the other hand, the recovery time is monotonically increasing with the parameter *χ*, and hence monotonically decreasing with the lysis rate *γ*.

## Discussion

Understanding bacterial responses to antibiotic treatments is critical for the future efficacy of antibiotic therapies. These responses are often studied at the level of single bacterial cells, which are not always predictive for the survival of bacterial populations due to cooperative interactions [22], or by using ill-defined measures of resistance, such as the minimal inhibitory concentration (MIC), which can be highly sensitive to variation in cell density and physiology [28, 54]. Cooperative interactions that enhance collective antibiotic survival can be caused by diverse mechanisms, in particular by the production of antibiotic-degrading enzymes, such as *β*-lactamases [22]. Given the substantial health problems caused by *β*-lactamase-producing infections [55], it is especially urgent to better understand the mechanisms and conditions promoting collective *β*-lactam survival.

Our present study was motivated by the observation of complex growth dynamics of two commonly used laboratory strains of *E. coli* in the presence of near-lethal concentrations of the cephalosporin cefotaxime (Figs. 1A and S3). To understand these dynamics, we performed experiments that identified the main causes of the observed sequence of growth, death and recovery of bacterial populations, including the initial biomass growth due to cell elongation (filamentation) and growth recovery allowed by the hydrolysis of cefotaxime by the chromosomally-encoded *β*-lactamase AmpC. Based on these key elements, we developed a kinetic model describing the growth dynamics of the two strains and validated it by comparing predicted and observed recovery dynamics (Fig. 5).

This work advances our current understanding of *β*-lactamase-mediated collective resistance in multiple ways. First, our model explicitly considers antibiotic hydrolysis at two distinct locations, in the periplasm of living cells and extracellularly after release of the enzyme by cell lysis [15, 43]. Both hydrolysis channels affect the extracellular antibiotic concentration in different ways, and our model allows to quantify their relative contributions to recovery of the population. As cell lysis progresses, the fraction of *β*-lactam that is hydrolysed extracellularly increases over time, and to what extent it contributes to recovery is therefore determined by the efficiency of hydrolysis via both channels and the amount of drug that needs to be removed (Fig. 6A). For the two *E. coli* strains considered in this study, our analysis shows that hydrolysis is primarily extracellular for REL606, but primarily periplasmic for K12. An immediate consequence of this result is that *β*-lactamase inhibitors with low periplasmic penetration [43] are expected to be more effective against REL606 than against K12.

Second, our model highlights two key properties of bacterial cells that determine the relative importance of periplasmic versus extracellular antibiotic degradation. One is the level of privatization of periplasmic hydrolysis of the antibiotic, represented by parameter *κ* (Eq. 2) and determined by the ratio of the total *β*-lactamase hydrolysis capacity of the cell and its permeability for the antibiotic. High values of *κ* correspond to high cell-level resistance and low contribution to antibiotic degradation in the environment and hence collective survival, and low values lead to larger relative contributions to environmental degradation at the expense of lower single-cell resistance. Therefore, variation in *κ* introduces a tradeoff between single-cell resistance and population-level resilience (see Fig. 6B). The other property is the lysis rate of the cell, represented by parameter *γ*, which contributes to extracellular degradation by releasing the *β*-lactamase. In experiments using an engineered strain with a fully private cytoplasmically-expressed *β*-lactamase with an inducible suicide module, Tanouchi et al. [56] found conditions where the death rate can be tuned to optimize collective survival. In our setting such an optimal lysis rate does not generally exist, as the recovery time is a weakly, but monotonically decreasing function of the lysis rate (Fig. 6B).

Third, it is worth noting that the higher lysis rate and higher degree of privatization that we inferred for *E. coli* REL606 conspire to yield similar values of the dimensionless parameter *η*, that quantifies the propensity for extracellular versus periplasmic CTX hydrolysis in the two strains. We can only speculate that this is indicative of a tradeoff that may exist more generally as a result of selection for population-level benefits. For example, cell death may be linked to *κ* via cell density-dependent induction of MazEF-mediated programmed cell death [57, 56], since bacteria with highly privatized *β*-lactamase function reach higher cell densities at a given antibiotic concentration due to their higher cell-level resistance. However, *γ* and *κ* may also be correlated as a passive consequence of the faster influx of antibiotic into the periplasm in bacteria maintaining a large gradient across their outermembrane (i.e. large *κ*), once periplasmic hydrolysis becomes insufficient to maintain periplasmic concentrations below the critical value.

Finally, our study highlights the special role of filamentation in *β*-lactamase-producing strains for collective survival. Filamentation is a known response to *β*-lactams via two pathways: as down-stream response of SOS induction [58], but also directly by binding of *β*-lactams like cefotaxime to PBP3, an essential component of the divisome [59, 60]. Filaments have been associated with enhanced *β*-lactam tolerance through delayed cell division [41, 61] and hence may build up larger biomass and *β*-lactamase concentrations before lysis sets in. Lysis rates of filaments have been shown to depend on the activation of cell division [60], as well as on filament length [34, 38, 42], which may allow for additional regulatory pathways to link *κ* and *γ* in addition to those described above. A more quantitative understanding of the physiological properties of filamentous cells seems required to fully understand their role in the evolution of resistance to *β*-lactams.

While we demonstrate potential collective benefits from the release of *β*-lactamase due to cell death in relatively homogeneous liquid bacterial cultures, these benefits may be enhanced in the structured environments, such as biofilms, in which bacteria typically live [18]. In contrast to other resistance mechanisms with only immediate survival benefits in the presence of the antibiotic, antibiotic-degrading enzymes like *β*-lactamases prime the environment in a way that regrowth is facilitated. This effect seems particularly important for fragmented populations in structured environments, where bacteria are cleared at some locations, which can subsequently be recolonized [62]. Another open question is how non-*β*-lactamase-producing community members living in close proximity to *β*-lactamase producers may benefit from this leaky public good [16], and how this has shaped the intra- and extracellular hydrolysis we observed within clonal populations. Clearly, these issues should be resolved to ensure the continued use of *β*-lactam antibiotics in combating infectious diseases.

## Acknowledgments

We are grateful to the Krug, de Visser, Bollenbach and Maier groups for inspiring discussions. We thank Leon Seeger for a thorough reading of the manuscript, and Gerrit Ansmann for help with statistical analyses. Further useful comments and insights were contributed by Eric Clément, Deniz Sezer, and the Kishony group. This work was funded by DFG SFB 1310 (German Research Foundation, Collaborative Research Centre 1310, project number 325931972).

## Contributions

R.G.: Conceptualization (equal); Investigation (lead); Methodology - experiments (lead); Writing – original draft (equal); Visualization (equal).

M.M.: Conceptualization (equal); Modeling and data curation (lead); Formal analysis (lead); Methodology - computational (lead); Writing – original draft (equal); Visualization (equal).

S.G.D.: Conceptualization (supporting); Writing – review (supporting).

M.Y.: Writing – review (supporting); Methodology - microfluidic experiment (supporting) and WGS (lead).

B.M.: Conceptualization (supporting); Writing – review (supporting).

T.B.: Conceptualization (supporting); Writing – review (supporting).

J.A.G.M.d.V: Conceptualization (equal); Project administration (equal); Writing – review and editing (lead); Funding acquisition (equal); Supervision (equal); Visualization (equal); Methodology - experiments (supporting).

J.K.: Conceptualization (equal); Project administration (equal); Writing – review and editing (lead); Funding acquisition (equal); Supervision (equal); Visualization (equal); Methodology - computational (supporting).

## Materials and methods

### Strains and media

The following ancestral strains were used in this study:

- *E. coli* REL606 knockout strain Δ*pcnB*.
- *E. coli* K12 Δ*lacA* strain JW0333 and Δ*ampC* strain JW4111 from the Keio collection [46].
- *E. coli* K12 strains from the ASKA collection that contain specific *amp*-genes on an IPTG inducible high copy number plasmid [45]. The inducible expression from the plasmid occurs in addition to the chromosomal expression, which is likely to lead to strong overexpression. The specific strains used were JW4111 (*ampC*), JW0106 (*ampD*), JW0107 (*ampE*), JW0423 (*ampG*) and JW5052 (*ampH*).

The two K12 strain groups encode resistance genes to chloramphenicol (ASKA) and kanamycin (Keio) that help to prevent contamination. The growth medium used in this study was Sigma-Aldrich LB (Lennox, L3022). Cefotaxime (Sigma-Aldrich, 219504) solutions were freshly prepared (see Supplementary Table 1 for concentrations used).

### Growth of *E. coli* strains REL606 and K12 in a linear CTX gradient

We inoculated 24 replicates cultures (inoculum size of 5 · 10^5^ cells per ml) in LB with a linear CTX concentration gradient of 40 CTX concentrations, ranging between 0.0 – 0.329 *µ*g/ml for Fig 1 and 0.0 – 0.113 *µ*g/ml for Fig S3. Each set of plates was arranged such that each plate contained four CTX concentrations from the gradient (two rows per concentration), leading to ten plates in total for the 24 replicates. Next, the 10 plates were loaded into a robot system (Liconic incubator at Prime robot) and optical density (OD) at 600 nm was measured approximately every 20 minutes over 60h in the case of Fig 1 and 24h in the case of Fig S3.

### Whole genome sequencing after recovery

To check for possible mutations causing the recovery of growth, genomic DNA was isolated from cell suspensions grown overnight at 0 and 0.08 *µ*g CTX/ml, using the NEB Monarch Genomic DNA Purification Kit (Ipswich, USA) according to the manufacturer’s instructions. A small aliquot of the isolated DNA was run on a 0.7% agarose gel with a 1 kb plus DNA Ladder (Thermo Scientific) to check for degradation. Non-degraded samples were sent to the Cologne Center for Genomics (Cologne, Germany) for NGS. Sequencing was performed on an Illumina Novaseq 6000 system with 150 bp paired-end reads and an average depth of 600X. Sequence analysis was performed using the pipeline described in [63, 64]. In short, the sequencing reads of the samples were mapped to the genome sequence of REL606 (NCBI Genbank RefSeq NC 012967.1, [65] using Burrows-Wheeler Aligner (v.0.7.17) [66]. Mapped reads were sorted with the sort function of samtools (samtools 1.8) and variants were called with the mpileup function and the call function from bcftools (bcftools 1.8) [67]. To identify deletions and duplications, genome coverage per base was calculated. After smoothing, the coverage with a sliding window of 30 bp, all segments in which the coverage dropped to zero for at least ten bases in a row were defined to be deletions. To be accepted as duplication, the coverage has to hit twice the genome-wide average coverage for ten subsequent bases. While we detected several mutations in both samples relative to the published genome of REL606, no mutations of any type were detected that distinguished the overnight cultures grown at 0 or 0.08 *µ*g CTX/ml, thereby ruling out mutations as cause for growth recovery.

### Cell growth in the microfluidic chamber

To track the growth behavior of individual bacteria over time, we used a microfluidic chamber. The flow chamber was made from polyoxymethylene with a cut for the main channel of size 6 *×* 56 *×* 1 mm with a hole in the center of size of 6 *×* 26 mm and two small holes at each side of the edge. The main channel was filled by polydimethylsiloxane (PDMS), forming a layer with a layout of 0.5 *µ*m high pillars and a 2 mm gap between each one of them. The two tubes with smaller holes allow media to flow through the chamber (further details of the chamber are given in [68]). To be able to track single cells over time, ~ 1, 000 cells were inoculated between a glass cover slide and a 1.5 mm-thick agar pad made of Milli-Q Water with 1% Bacto agar (BD). This was integrated into the flow chamber and sealed by picodent twinsil (Picodent) to restrict the flow of the media between the agar pad and the PDMS, and to prevent leaks. After the flow chamber was sealed, it was placed on the stage of a 30^°^C heat-controlled inverted microscope (Nikon Eclipse TE2000-E) using a 100X objective. Once the field of view for the imaging was chosen, Differential Interference Contrast (DIC) images were taken every 10 minutes. Two syringes containing media were connected consecutively to the fluidic chamber and controlled by a syringe pump.

The experiment was done in two steps: first, only the syringe with medium containing nutrients and 0.2 *µ*g CTX/ml was opened at a flow rate of 60 *µ*l/minute for 2.5h. In the second step, only the syringe with CTX-free nutrient medium was opened at a flow rate of 15 *µ*l/minute for 20h.

### Spent medium experiments

To investigate the antibiotic degradation by the cells, we performed three spent medium experiments. In the first experiment, we aimed to determine whether protection was due to CTX degradation in the spent medium or goes beyond that. In the second experiment, we quantified the level of protection offered by the spent medium. In the third experiment, we tested whether protection was due to a protein and sensitive to heating. For all three experiments, we prepared culture media by mixing the spent medium with different amounts of CTX and fresh nutrients.

To generate the spent medium, we grew *E. coli* in LB containing 0.08 *µ*g CTX/ml, which induces non-monotonic growth behaviour. The incubation time was 14h, corresponding to the time point at which recovery occurs. After incubation, we filtered out the cells to obtain the spent medium. In the second experiment, we additionally sonicated the medium for 10 seconds before filtration to remove effects from variation in cell lysis during preparation.

In order to obtain a mixed culture medium with a total volume of *V* = 200 *µ*l, we mixed three different media:

- A bacterial culture with a culture volume *V*_*b*_ = 20 *µ*l: freshly made LB without CTX was prepared with a cell density of 5 · 10^6^ cells per ml. In the total culture volume *V* this gives a final inoculum size of 5 · 10^5^ cells per ml.
- Diluted spent medium with a culture volume *V*_*s*_ = 90 *µ*l: The harvested spent medium was mixed in different proportions with freshly made LB without CTX. The fractions of the spent medium in this volume were chosen to be 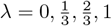, leading to the spent medium fractions 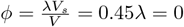, 0.15, 0.30, 0.45 in the total culture volume *V*.
- Fresh medium with [CTX] in a culture volume *V*_*f*_ = 90*µ*l: Freshly made LB was prepared with a linear CTX gradient. The CTX concentrations 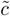 in this volume were chosen in such a way that, under the assumption that no CTX was left in the spent medium, the CTX concentration in the total culture volume *V* would amount to the prescribed nominal concentration [CTX]_*m*_. This implies that 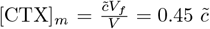, which determines 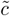 for a given nominal concentration.

The spent medium fractions and CTX concentration used in the different experiments were chosen as follows:

- Experiment 1: Bacterial cultures with spent medium fractions of *ϕ* = 0% and 45% and fresh medium containing CTX at concentrations 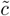 ranging from 0 to 1.222 *µ*g/ml in steps of 0.111 *µ*g/ml, leading to nominal concentrations from 0 to 0.55 *µ*g/ml in steps of 0.05 *µ*g/ml.
- Experiment 2: Bacterial cultures with spent medium fractions of *ϕ* = 0%, 15%, 30%, 45% and fresh medium containing CTX at concentrations of 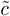 ranging from 0 to 0.777 *µ*g/ml in steps of 0.111 *µ*g/ml, leading to nominal concentrations from 0 to 0.35 *µ*g/ml in steps of 0.05 *µ*g/ml.
- Experiment 3: Bacterial cultures with spent medium fractions of *ϕ* = 0% and 45%. For both fractions, the medium was split into two portions, one of which was heated to 98^°^C for 30 minutes. Subsequently, the heated and non-heated media were mixed with fresh medium containing CTX at concentrations of 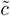 ranging from 0 to 0.777 *µ*g CTX/ml in steps of 0.111 *µ*g CTX/ml, leading to nominal concentrations ranging from 0 to 0.35 *µ*g CTX/ml in steps of 0.05 *µ*g CTX/ml.

After the preparation of the culture medium for each experiment, we measured the OD growth curves using a plate reader.

### Protection ability of spent medium in a structured environment

We poured six plates with 20 ml 2% LB + 1.5% agar and 0.25 *µ*g CTX/ml. After the agar in the plates solidified, we placed a drop of 200 *µ*l of spent medium (harvested after 14 h of growth in LB with 0.08 *µ*g CTX/ml) on three different plates, and a drop of 200 *µ*l of fresh LB without CTX on the three other plates (marked by red circles in Fig. S2). Then we let the drops be soaked into the medium for 6h. After the plates were dry, we inoculated a drop of 100 *µ*l of *E. coli* (5 · 10^5^ cells) and distributed them all over the plate with bacterial cell spreaders. Finally, we incubated the plates overnight at 30^°^C degrees and subsequently imaged the plates.

### Protection ability of spent medium from ASKA and Keio strains

*E. coli* JW4111 was grown with inoculum size of 5 · 10^5^ cells/ml in three different concentrations of IPTG (0, 0.1 and 1 mM) for 24 h without CTX, cells were filtered out, and the resulting spent medium was mixed with fresh medium (to a final spent medium fraction of 45%) containing a different amount of CTX (see Exp. 7 in Table S1) and OD of the cultures was measured over time.

We repeated the experiment at intermediate AmpC induction, growing *E. coli* JW4111 with inoculum size of 5 · 10^5^ cells/ml with 0.1 mM IPTG overnight and harvesting the spent medium. The spent medium was then split into two fractions and 5*µ*g/ml of boronic acid (a *β*-lactamase inhibitor) was added to one fraction.

We performed two further experiments. First, we over-expressed (by using 1 mM IPTG) all *amp* genes separately using strains from the ASKA collection (JW4111 (*ampC*), JW0106 (*ampD*), JW0107 (*ampE*), JW0423 *ampG*) and JW5052 (*ampH*)) and measured their growth in varying CTX concentrations (See Exp. 14 in Table S1). Second, we grew ΔampC and ΔlacA strains from the KEIO deletion library overnight in LB broth without CTX. Then both cultures were subjected to sonication separately for 2 minutes to lyse all cells, and the spent media were harvested. Each harvested medium was mixed with fresh medium (to a final spent medium fraction of 45%) with a CTX gradient (See Exp. 9 in Table S1). The OD growth of *E. coli* K12 ΔlacA in each of the mixed media was then measured.

### Nitrocefin assay of *β*-lactamase activity in the absence of CTX

Nitrocefin is a chromogenic cephalosporin whose hydrolysis rate can be measured to quantify the *β*-lactamase activity of a culture. Here, we measured *β*-lactamase activity of both *E. coli* strains in the absence of CTX to assess differences in per-cell AmpC expression level under the assumption that K12’s and REL606’s AmpC’s have similar CTX hydrolysis capacity.

We made a nitrocefin solution stock by dissolving 5 mg nitrocefin in 500 *µ*l dimethyl sulfoxide (DMSO, Sigma Aldrich) and added 9.5 ml phosphate buffer (100 mM, pH 7) to obtain a total volume of 10 ml. The stock solution was kept in a freezer at − 20^°^C for a maximum of 2 months. We performed nitrocefin assays on the spent medium collected from overnight cultures of both strains to quantify the amount of *β*-lactamase produced by the cells in the absence of antibiotic stress. We grew eight replicate cultures for both strains (REL606 and K12) for 22 h in LB without CTX, and subsequently split each into two subpopulations. One subpopulation was subjected to sonication for 2 minutes to lyse all cells, and the medium was filtered (using a a sterile 0.2 *µ*m syringe filter) to harvest the sonicated spent medium. The other subpopulation was centrifuged (12k RPM for 10 minutes), after which the cells were filtered out with a sterile 0.2 *µ*m syringe filter, to obtain the not-sonicated spent medium. Then we mixed 100 *µ*l of the spent media with 100 *µ*l of the 10 *µ*g/ml nitrocefin solution stock, diluting the nitrocefin solution stock concentration by a factor of 50, in a 96-well plate. The *β*-lactamase activity was quantified by the rate of change in the red absorption (OD490) due to nitrocefin hydrolysation (https://www.sigmaaldrich.com/deepweb/assets/sigmaaldrich/product/documents/400/065/mak221-tech-bulletin-mk.pdf and [23]).

## Supplementary Table

**Table S1:** The table lists the CTX concentration ranges used to obtain the experimental results presented in the corresponding figures.

| Experiment | Figure | [CTX] |
| --- | --- | --- |
| Exp 1 | Fig 1 a | 0 – 0.329 $\mu\text{g/ml}$ |
| Exp 2 | Fig 1 c | 0.08 $\mu\text{g/ml}$ |
| Exp 3 | Fig 2 | 0.2 $\mu\text{g/ml}$ |
| Exp 4 | Fig 3 a | 0 – 1.222 in steps of 0.111 $\mu\text{g/ml}$ |
| Exp 5 | Fig 3 b | 0 – 0.777 in steps of 0.111 $\mu\text{g/ml}$ |
| Exp 6 | Fig 3 c and d | 0 – 0.777 in steps of 0.111 $\mu\text{g/ml}$ |
| Exp 7 | Fig 4 a | 0.006 – 0.113 in steps of 0.003 $\mu\text{g/ml}$ |
| Exp 8 | Fig 4 b | 0.047 – 0.398 in steps of 0.035 $\mu\text{g/ml}$ |
| Exp 9 | Fig 4 c | 8-fold increases between 0.008 and 5.625 $\mu\text{g/ml}$ |
| Exp 10 | Fig S1 a | 0.094 $\mu\text{g/ml}$ |
| Exp 11 | Fig S1 b | 0.038 – 0.206 in steps of 0.028 $\mu\text{g/ml}$ |
| Exp 12 | Fig S2 | 0.08 and 0.25 $\mu\text{g/ml}$ |
| Exp 13 | Fig S3 | 0 – 0.113 $\mu\text{g/ml}$ |
| Exp 14 | Fig S4 | 4-fold increases between 0.002 and 10 $\mu\text{g/ml}$ |

## Supplementary figures

**Figure S1:**
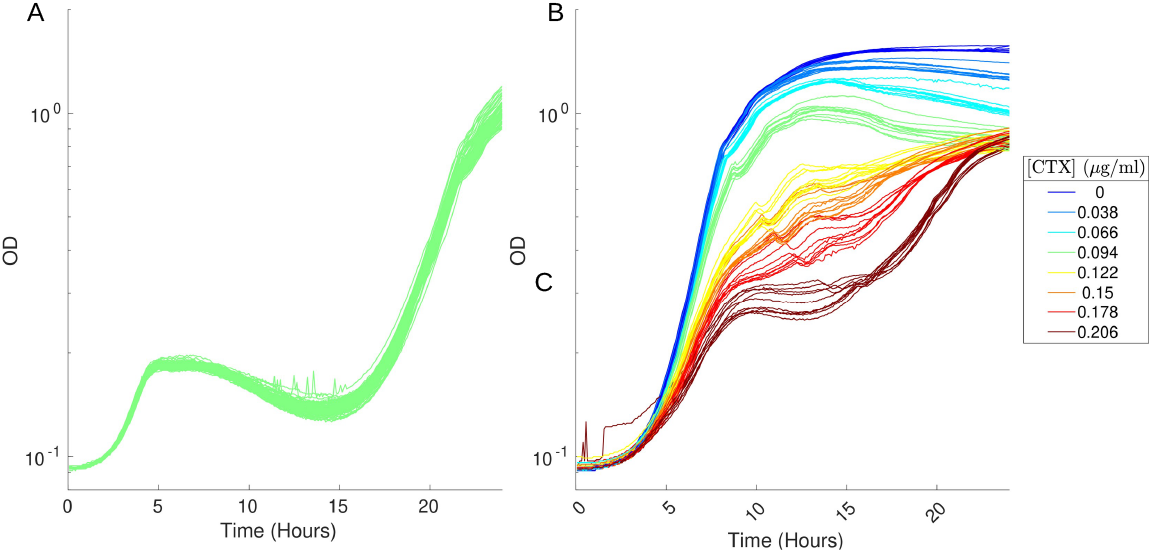
Recovery growth is not due to resistance mutations. The experiment was done in two phases. **A**: In the first phase, 80 replicate cultures of *E. coli* REL606 were grown at 0.094 *µ*g CTX/ml for 24h. **B**: We then harvested the cells, washed them with fresh LB medium, and created 10 independent cultures by pooling eight replicate cultures for each inoculum. Dilutions from these inocula were used to start 80 new cultures in the presence of eight CTX concentrations.

**Figure S2:**
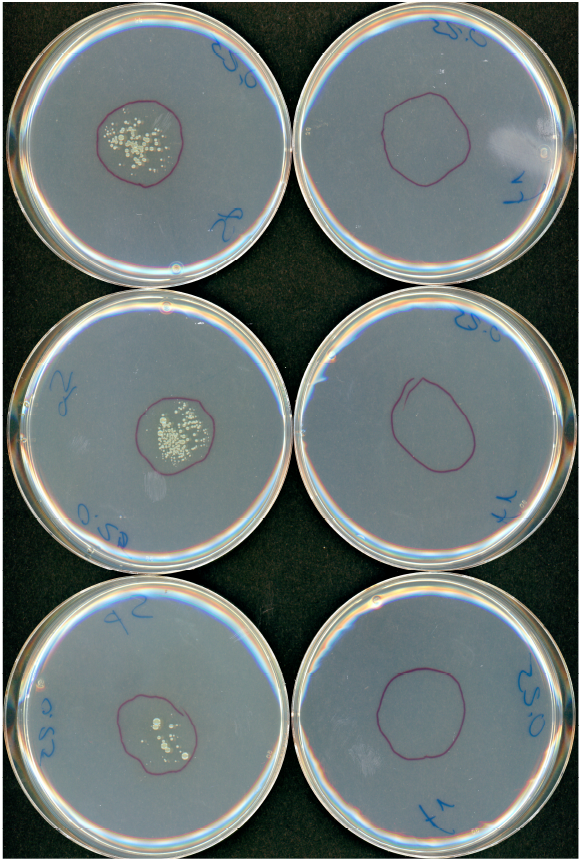
Test of protection by spent medium on agar. The figure shows six plates prepared with 20 ml LB with agar and 0.25 *µ*g CTX/ml. In the left column, the red circles represent the area where 200 *µ*l drops of spent medium from cultures grown for 14 h in the presence of 0.08 *µ*g CTX/ml were placed and allowed to dry for 6 h. In the right column, the circles represent the area of control drops of fresh medium. Visible growth is detected only in the area covered by spent medium.

**Figure S3:**
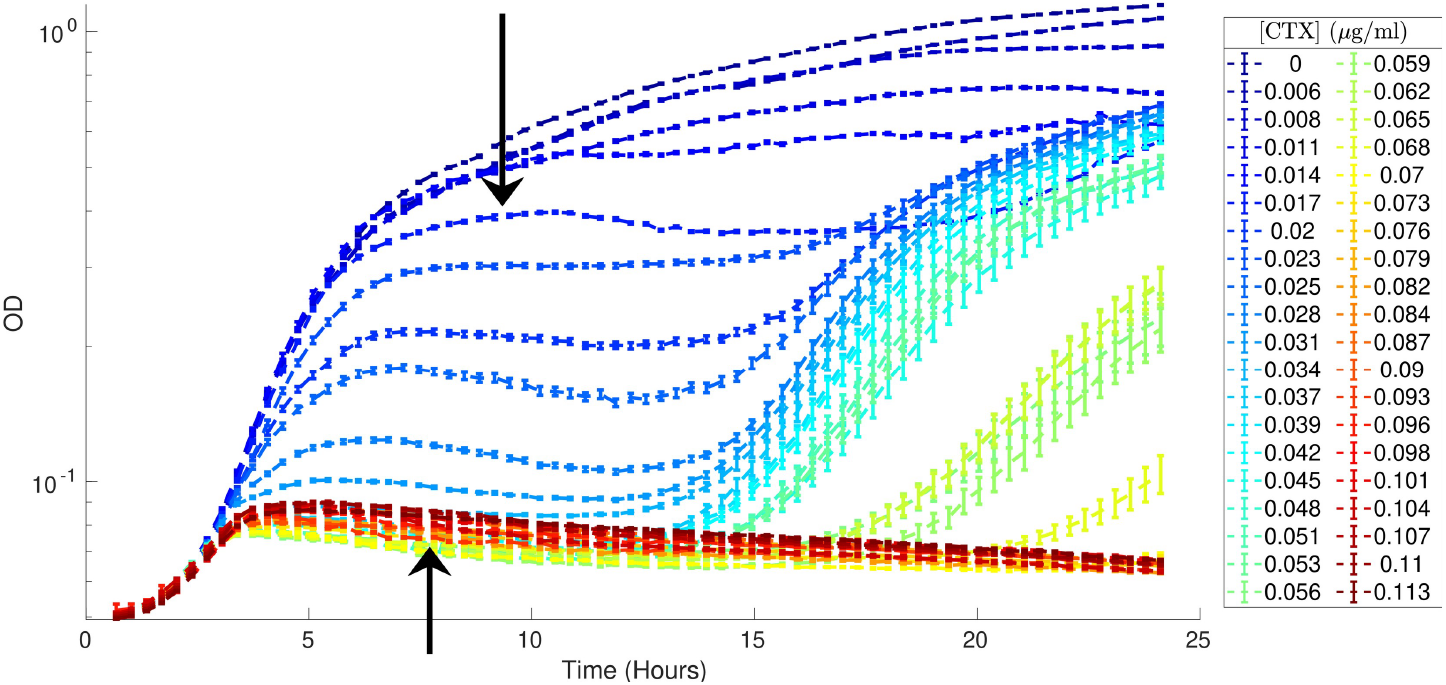
Growth curves of *E. coli* K12 in a linear CTX concentration gradient. Shown are the average and SD of OD obtained from 24 cultures of *E. coli* K12 in a linear CTX concentration gradient, ranging from 0 – 0.113 *µ*g/ml. The inoculum size was 5 · 10^5^ cells/ml (see Materials and Methods for details). For concentrations below 0.014 *µ*g CTX/ml, the growth curves increase monotonically, saturating at late times. For concentrations above 0.014 *µ*g CTX/ml and below 0.076 *µ*g CTX/ml (indicated by the arrows), the growth curves exhibit non-monotonic behavior: initial growth leads to a maximum, which is followed by a decay of biomass until approximately *t* = 12 h, before growth resumes. For CTX concentrations above 0.076 *µ*g/ml, the initial growth is followed by monotonic biomass decay.

**Figure S4:**
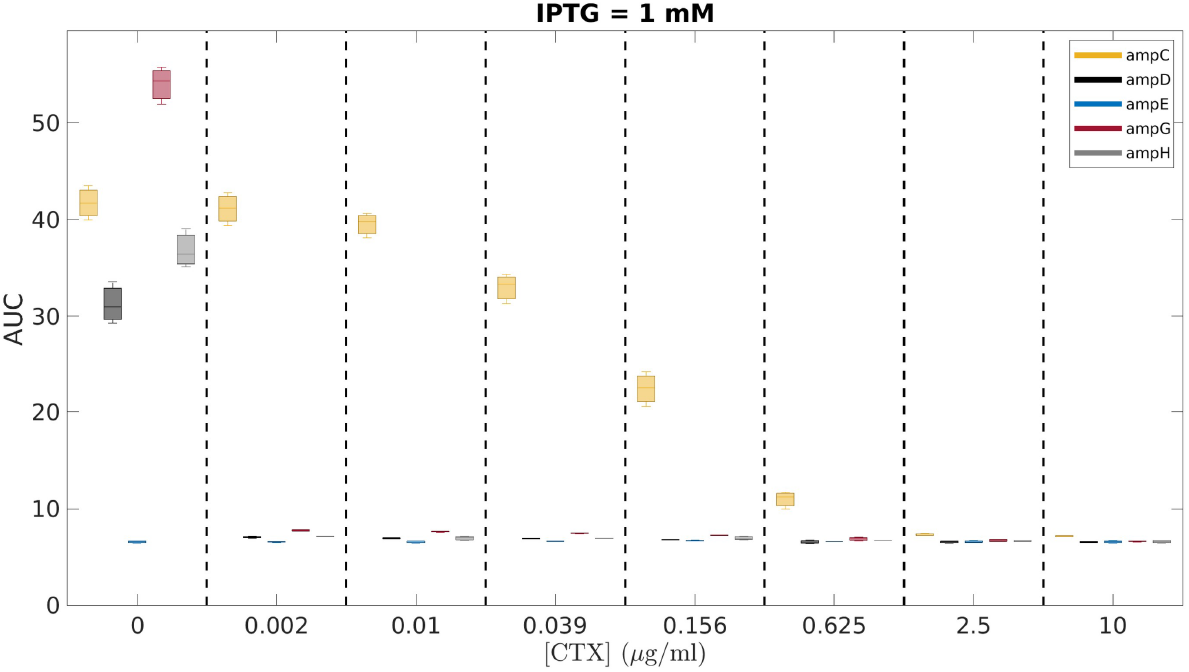
*AmpC* is the only *amp* gene whose expression increases CTX resistance. The figure shows the area under the OD-curves (AUC) of the first 30 h of growth for five different ASKA strains, in which the genes *ampC, ampD, ampE, ampG*, and *ampH* were overexpressed at an IPTG inducer concentration of 1 mM. Each bin corresponds to one CTX concentration.

## Supplementary Information: Data analysis and modeling

### Feature extraction from OD curves

Figure S5 shows a typical family of 40 OD curves obtained from the growth of replicate cultures. Each OD curve represents growth under a different concentration of CTX and is color-coded from light orange to dark red, as indicated in the legend of the figure. For low concentrations of CTX, the OD growth is monotonic with saturation at late times.

**Figure S5:**
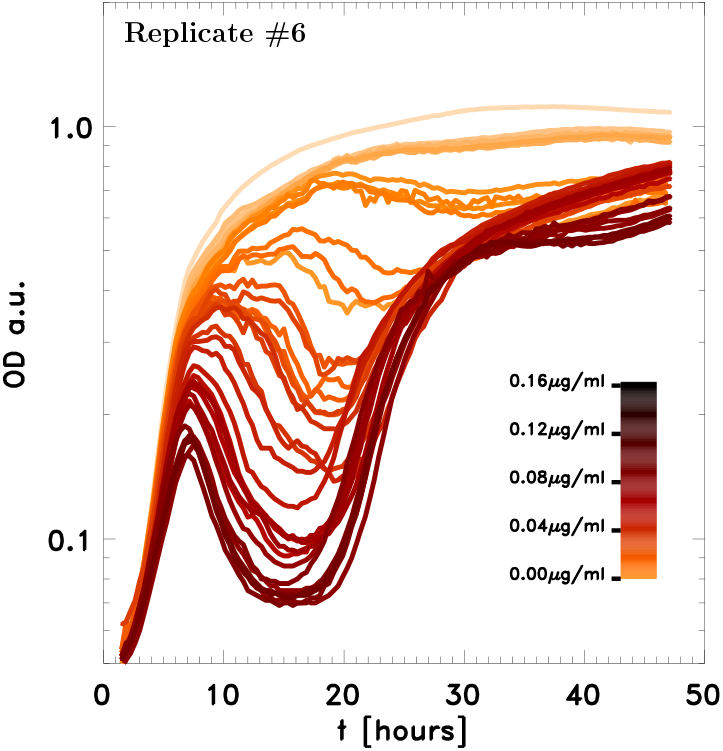
Family of OD growth curves of the REL606 strain in a linear gradient of CTX concentrations, color coded as shown in the legend. With increasing concentrations, the curves cross over from monotonic to non-monotonic growth.

We characterize the growth curves by the time *t*_*G*_ at which the exponential growth rate is maximum. As the CTX concentration is increased, the monotonic OD growth crosses over into non-monotonic growth. Figure S6 shows a sample non-monotonic OD curve for *E. coli* REL606 at a CTX concentration of 0.085 *µ*g/ml. The curve exhibits a local maximum and minimum of the OD at the points labeled *P* and *Q*, respectively. These points mark three phases of population growth: an initial growth phase characterized by a maximum growth rate at point *G*, an intermediate decay with the decay rate becoming largest at *L*, and finally a recovery growth phase with maximum recovery growth attained at point *R*.

We implemented an automatic work flow to extract these features from the measured OD curves, determining first whether the growth is monotonic or not. If so, we identify the point *G*, as well as the time and corresponding exponential growth rate at *G*. If the growth is non-monotonic, we also determine the maximum decay – and if present – recovery growth rates, the times where these are attained, as well as the times and OD values of the local extrema, points *P* and *Q*. In the following we provide details on the extraction of these features.

We estimate instantaneous growth rates by log-fitting a moving window of *n*_fit_ consecutive OD data points to a straight line. Note that OD values are measured every 20 minutes. Since the initial growth rate attains its maximum in a rather short time (less than 8h after the start of the experiment), we chose *n*_fit_ = 5 data points for the window width, corresponding to a time window of 100 minutes. For the estimation of all other growth rates, we chose *n*_fit_ = 12. We defined the growth rate as the maximum of the log-fitted instantaneous rates over a time interval from *t*_min_ = 1.2*h*h to 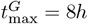, as indicated by the corresponding dashed vertical lines in Fig. S6.

For the determination of the intermediate decay and recovery growth rates and the local OD maxima and minima, we first check whether the OD curve is non-monotonic. We do this by testing for the existence of a local maximum of the OD. Since the OD curves, particularly around the transition from monotonic to non-monotonic growth, are noisy, we first pass the curves through a Gaussian filter in order to smoothen them. The thick curve in light grey shown in Fig. S6 is the filtered version of the measured OD curve shown in orange-red. We use the filtered OD curves to test for the existence of a local extremum. In practice, we do this by setting a maximum time 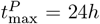, such that if a local OD maximum is not found before that, we regard the OD curve as monotonically increasing. If a maximum and hence the point *P* was identified, we also checked whether the OD curve reaches a local minimum afterwards. If so, we identify the OD value and time *t*_*Q*_ where it was attained.

**Figure S6:**
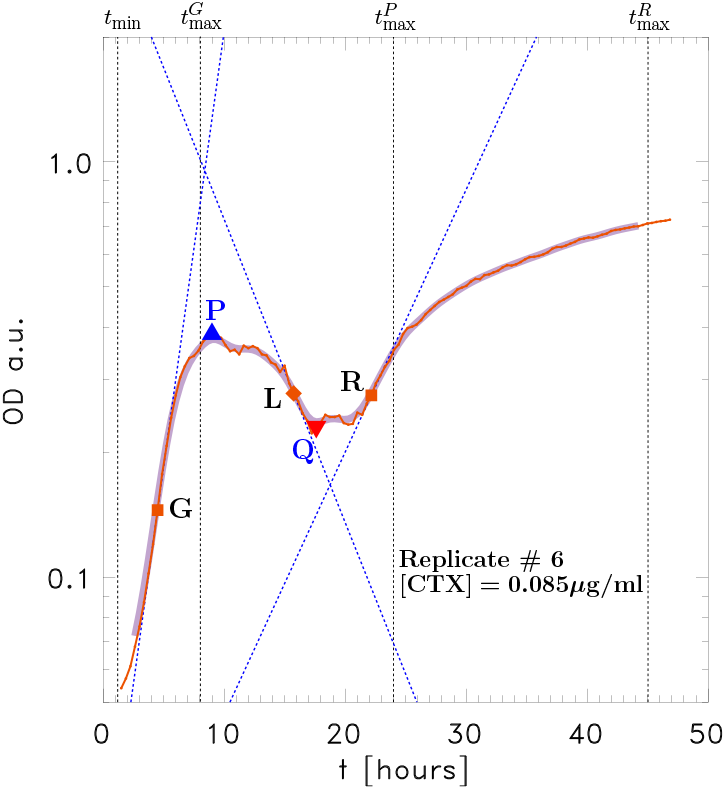
A sample non-monotonic OD curve, illustrating the features of interest that we extract from it. These are the points *P* and *Q* where the OD curve attains its local maximum and minimum, if they exist, as well as the points *G, L*, and *Q*, where the rates of initial OD growth, intermediate decay, and recovery are largest. The times 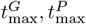, and 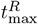 are the cut-off times up to which we search for these features, and are indicated by the corresponding dashed vertical lines. Refer to the text for further details.

In the case that the OD curve is non-monotonic, we also determine the point *L* where the intermediate decay rate is maximum, which must be located in time after *t*_*P*_. If a local minimum *Q* was found, we search for the maximum of the recovery growth rate in the time interval starting from *t*_*Q*_ up to a cut-off time 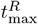. For the REL606 data, for which time-series were obtained up to *t* = 60h, we used an upper cut-off of 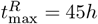, while for the K12 data the cut-off was 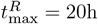.

The dashed blue lines in Fig. S6 indicate the fitted OD growth rates. These are the data points plotted in Fig. 1B of the main text. As noted before, for REL606 and at experimental observation times larger than *t* = 45h, the recovery of cultures subjected to high CTX concentrations shows considerable fluctuations from replicate to replicate, as evidenced by the large standard deviations in Fig. 1A. We, therefore, discarded the experimentally observed recovery rates for CTX concentrations larger than 0.250 *µ*g/ml.

### Modeling CTX hydrolysis in the periplasm and extracellular medium

Our modeling approach involves linking the enzyme production, i.e. via cell growth and its subsequent release into the extracellular medium by lysis, with the known kinetics of CTX hydrolysis by *β*-lactamases. Similar modeling approaches have been put forward before. Andreani and collaborators have considered hydrolysis due to cell lysis and subsequent release of *β*-lactamases into the extracellular medium [34, 42]. They assume that lysis starts to occur once the filamented cell length exceeds a CTX concentration dependent critical length. This length marks the onset where the rate at which damage to the peptidoglycan layer due to CTX exceeds the rate at which the cell can repair the damage. Artemova and collaborators analysed a model where CTX is hydrolysed both inside the periplasm and in the extracellular medium. Lysis is assumed to occur once the CTX concentration inside the periplasm exceeds a threshold value, which they call the single cell MIC [15]. Recently, Geyrhofer and collaborators have considered a model that does not explicitly take lysis into account, but assumes that there is an outflux of *β*-lactamase into the extracellular medium from intact cells [28].

Our modeling strategy follows that of Artemova et al. [15], but in addition we estimate the production rates of viable and lysed biomass from the empirical OD curves that we obtained at various CTX concentrations. We use these rates as inputs to our modeling of the chemical kinetics of CTX hydrolysis by *β*-lactamases. We further assume that the *β*-lactamases are confined to the periplasm where they can diffuse freely, unless the cells lyse [69]. In our model, hydrolysis of CTX by *β*-lactamase can therefore take place either inside the periplasm of non-lysed cells or in the extracellular medium, in which case the amount of *β*-lactamases available at any given time will depend on the biomass that has lysed so far. We ignore processes that lead to a natural degradation of CTX or *β*-lactamases, assuming that the corresponding decay times are much larger than the duration of our experiments.

The diffusion of *β*-lactams across the outermembrane bilayer is negligible and their entry into the periplasm occurs through the porins [70, 71]. The influx of CTX via porins into the periplasm is assumed to be governed by Fick’s law

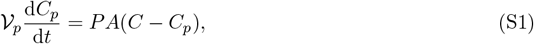

where *P* is the permeability constant, *A* is the *E. coli* surface area, 𝒱_*p*_ is the periplasmic volume, and *C* and *C*_*p*_ are the CTX concentration outside the cell and inside the periplasm, respectively.

The hydrolysis of CTX via *β*-lactamases in the periplasm follows a Michaelis-Menten type kinetics:

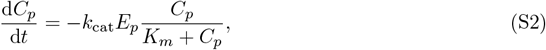

where *E*_*p*_ is the concentration of *β*-lactamase, while *k*_cat_ and *K*_*m*_ are the rate constants of the CTX hydrolysis reaction kinetics.

An equilibrium is established once the concentration of CTX inside the periplasm is constant in time, implying that the influx rate of CTX via porins is balanced by the hydrolysis rate inside the periplasm, leading to the Zimmermann-Rosselet Equation [51]

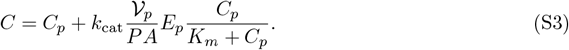

It is convenient to go to units in which the concentrations are measured relative to *K*_*m*_. Thus defining *a* = *C/K*_*m*_ and *a*_*p*_ = *C*_*p*_*/K*_*m*_ the influx and hydrolysis equations are given by

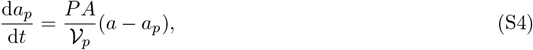

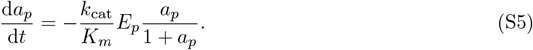

Thus (S3) becomes

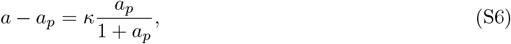

where we have introduced the dimensionless quantity

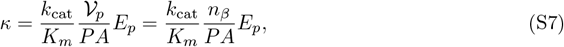

which essentially is the ratio of the rate of CTX hydrolysis to the influx rate. Here *n*_*β*_ = 𝒱_*p*_*E*_*p*_ denotes the number of *beta*-lactamase molecules in a normal-size cell.

It is clear that the CTX gradient *a* − *a*_*p*_ across the outermembrane depends on the value of *κ*. If *κ* ≪ 1, then the hydrolysis rate is small compared to the influx rate and from (S6) we see that *a* − *a*_*p*_ is of the order of *κ*. Hence the difference of CTX concentrations outside and inside the periplasm is small. Conversely, if *κ* ≫ *a*, then the hydrolysis rate is relatively high, meaning that any CTX molecule that diffuses into the periplasm is rapidly hydrolysed so that large concentration differences can be maintained. Indeed we find that in this limit, *a/a*_*p*_ is of order *κ*.

For *E. coli* K12, Nikaido and collaborators have measured the permeability *P*, as well as the reaction constants for CTX hydrolysation by the chromosomally encoded *β*-lactamase AmpC, finding *P* = 1.8 · 10^−5^ *cm/s, k*_cat_ = 0.06 *s*^−1^ and *K*_*m*_ = 0.16 *µM* [48, 49]. Assuming an *E. coli* volume of 1.3 *µm*^3^, a surface area of 6.28 *µm*^2^, and estimating the volume of the periplasm to be 30% of the cell volume [72], we find that *κ* depends on the number *n*_*β*_ of AmpC molecules per cell volume as *κ*_*K*12_ = 5.5 · 10^−4^ *n*_*β*_. Typical values for *n*_*β*_ are of the order of a few hundreds, and depend on the composition of the culture medium [52, 53] (see also https://biocyc.org/gene?orgid=ECOLI&id=EG10040#showAll).

**Figure S7:**
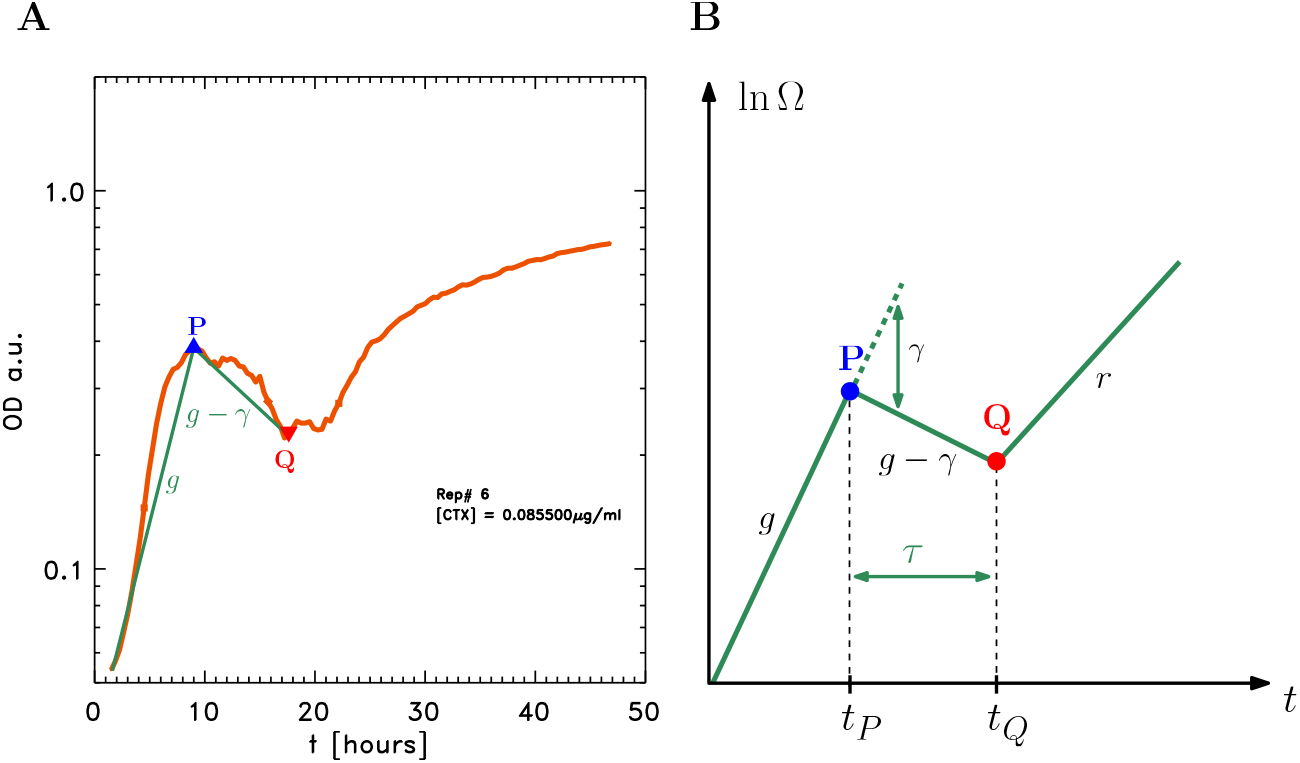
Modeling *E. coli* growth. **A**: Experimentally obtained OD curve for *E. coli* strain REL606 at a CTX concentration of 0.0855 *µ*g/ml. The OD reaches a peak OD_*P*_ at time *t*_*P*_ that is followed by a decay and subsequent onset of recovery at *t*_*Q*_ and OD_*Q*_. The green lines represent an average constant initial growth and intermediate decay rate, as inferred from the OD curve. **B**: The simplified growth curve used in our modeling. We assume initial growth of biomass at fixed exponential growth rate *g*, which is followed at time *t*_*P*_ by simultaneous decay due to lysis and continuing growth, given by the effective rate *m* = *g* − *γ*. Here *γ* is the rate of cell lysis. At point *Q* exponential growth resumes again with some growth rate *r*. The recovery time *τ* = *t*_*Q*_ − *t*_*P*_ is defined to be the time between the onset of lysis and the beginning of recovery.

Another point worth stressing is that for the values quoted in the main text, it follows from (S4) and (S5) that the equilibration of the CTX concentration inside the periplasm is very rapid and reached within seconds [24, 48]. Since in typical settings, such as our experiments, the CTX is hydrolysed over a time scale of hours, this means that the CTX concentration inside the periplasm is in quasistatic equilibrium throughout, i.e we can assume that (S6) holds although *a*_*p*_ and *a* change (sufficiently slowly) in time. We will refer to this as the quasistatic approximation.

For future reference, let us note that (S6) can be inverted to yield *a*_*p*_ as a function of *a* and *κ* as

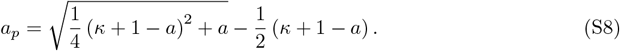

### Coupling CTX hydrolysis to *E. coli* growth and lysis

We are interested in modeling the non-monotonic growth and recovery of the population at intermediate CTX concentrations. Figure S7A shows the typical OD behavior. Initially, the OD grows, reaching local maximum OD_*P*_ at point *P* and time *t*_*P*_. This point marks the onset of OD decay continuing up to point *Q*, which is reached at time *t*_*Q*_ and OD_*Q*_. For times larger than *t*_*Q*_, the OD starts to grow again, which marks the onset of recovery of the bacterial population. Panel B is a simplified picture of this OD behavior, expressed now in terms of the biomass Ω. We assume that up to time *t*_*P*_ the biomass growth is exponential with fixed growth rate *g*, while between *P* and *Q* we assume that the biomass is decaying at a rate *m* = *g* − *γ*, where *γ* is the cell lysis rate, assumed to be constant in time. Beyond *Q*, the biomass resumes exponential growth with rate *r*.

In the present section, we will assume that the growth of biomass, as shown in Fig. S7B, is given as input into our modeling and specified in terms of the biomass Ω_*P*_ at *P* and time *t*_*P*_, as well as the *mean* initial growth and intermediate decay rates *g* and *m*. For the given range of CTX concentrations, we can estimate these parameters from the measured OD curves, as indicated by the green lines in panel A of Fig. S7. Assuming that recovery sets in once the CTX concentration in the medium has fallen due to hydrolysis below a certain concentration, our goal is to work out the time when this occurs, i.e. the recovery time *τ* = *t*_*Q*_ − *t*_*P*_.

Let us emphasize that we assume that the growth and subsequent lysis rates are constant in time, but that their overall values can depend on the antibiotic concentration. However, the antibiotic concentration in turn does depend on time, due to the hydrolysis. In our modeling, we will consider this time dependence of growth rates as a second order effect, which we take care of by assuming that *g* and *γ* denote effective or mean growth and lysis rates, that thereby take into account in part an implicit time-dependence due to the change of concentrations. Practically, this is conveniently achieved by approximating the curved pieces of the log OD curve by linear segments, which leads to the piece-wise linear growth of log-biomass as shown in Fig. S7A and B. Following Geyrhofer et al. [28], it is possible to incorporate an explicit dependence of growth and lysis rates on instantaneous antibiotic concentration. While such a modeling is more realistic and will also round out the biomass growth at points *P* and *Q*, it comes at the price of analytical complexity.

Since our main aim of modeling is to obtain a qualitative as well as rough order of magnitude quantitative understanding, we prefer to work with effective and constant exponential growth and decay rates. Figure S8A and B show the dependence of the effective growth rate *g* and decay rate *g* − *γ* on the initial CTX concentration [CTX]_ini_, obtained from the OD curves for *E. coli* K12 and REL606. These values are used as input for our numerical results shown in Fig. 5 of the main text, as well panels C and D of Fig. S8.

**Figure S8:**
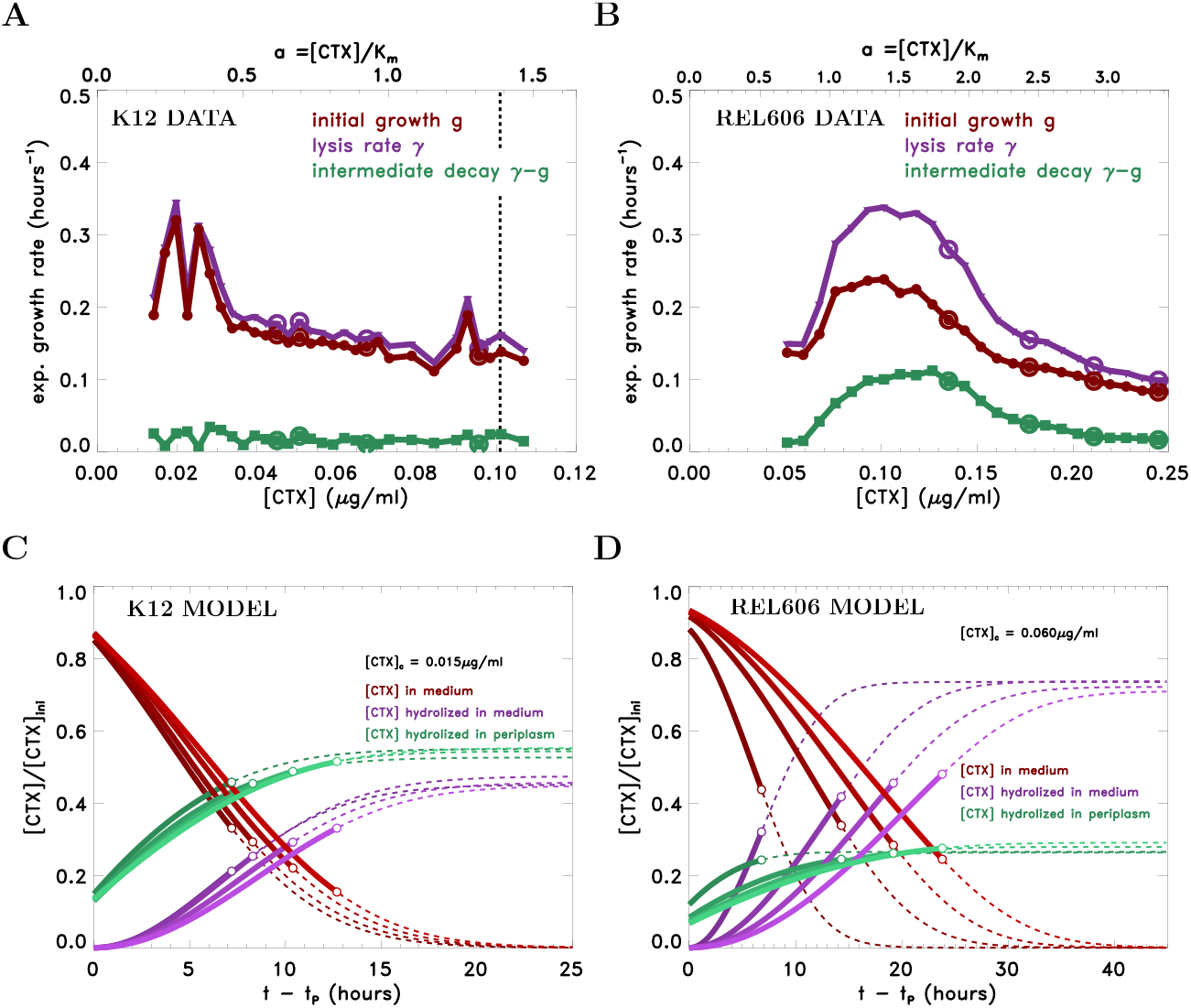
Data-supported model predicting the amounts of CTX hydrolysed by AmpC in the medium and the periplasm. **A** and **B:** Empirical dose-response curves depicting the average empirical growth, decay, and lysis rates as function of initial CTX concentration for *E. coli* K12 and *E. coli* REL606. The encircled data points show the concentration values for which these rates were used as inputs for the model predictions shown in panels C and D. **C** and **D:** Time evolution of the CTX concentration (red curves) predicted by the model for a range of concentrations, using the empirical biomass growth and lysis rates. The range of initial concentrations is color coded with high and low values corresponding to dark and light shades. The curves in green and purple show the amount of CTX that has been hydrolysed in the periplasm and the medium, respectively.

For the range of CTX concentrations that give rise to non-monotonic OD growth, we assume that in the initial growth state cells filament, but do not lyse, so that cell length grows as

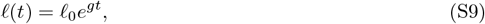

where *𝓁*_0_ is the length of a cell of normal size. Assuming an inoculum size of *N*_0_ cells, the (linear) biomass grows as

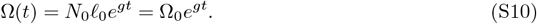

up to time *t* = *t*_*P*_. At *P*, cells start to lyse at a constant rate *γ*. Thus for *t > t*_*P*_, cells continue growing at exponential rate *g*, while some of them lyse at rate *γ* and we find

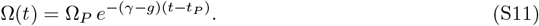

We first work out the reaction kinetics of CTX hydrolyzation outside the cells, i.e. in the extracellular medium. This process is driven by the release of *β*-lactamase molecules into the extracellular medium by lysis, leading to a gradual rise of the enzyme concentration. Let *E*_ly_(*t*) be the concentration of *β*-lactamase in the extracellular medium so that *E*_ly_(*t*) = 0 for *t* ≤ *t*_*P*_ (no lysis yet). For *t > t*_*P*_, we assume that the break-down of CTX is by hydrolysis and follows the Michaelis-Menten type reaction kinetics discussed before,

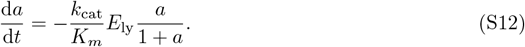

The amount of *E*_ly_(*t*) accumulates as a result of the lysis of cells and hence is given by

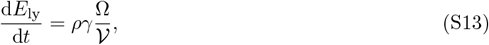

where *γ*Ω is the amount of biomass lysed per unit time, 𝒱 is the volume of the container, and *ρ* is the number of *β*-lactamase molecules per biomass. Since the time evolution of Ω is known, and setting *τ* = *t* − *t*_*P*_, we have

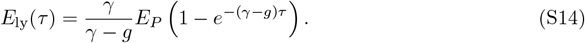

Here, *E*_*P*_ is the effective concentration of *β*-lactamase in the extra cellular medium that would have been attained if all the cells present at *P* had lysed at once. This concentration depends on the live biomass at *P* as well as the number of enzyme molecules per biomass, which in turn depend on the inoculum size.

Substituting (S14) into (S12), we find that the rate at which CTX is hydrolysed in the extra-cellular medium is given by

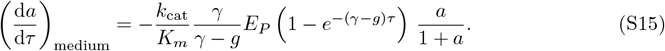

Let us now turn to the hydrolysis of CTX inside the periplasm. Recalling that we are in the quasistatic regime, the concentrations inside and outside the perisplam are related by (S6). The total flow rate of CTX into the cells by diffusion is given by

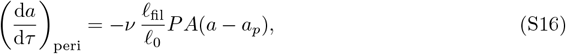

where *ν* is the concentration of cells, i.e. number of (filamented) cells per unit volume, while *A* and *A𝓁*_fil_*/𝓁*_0_ are the surface area of normal-size and filamented cells, respectively. Here *𝓁*_fil_ is the length of filamented cells at time *τ*, cf. (S9). At equilibrium, (S16) furnishes also the rate at which the CTX is hydrolysed inside the periplasm as follows. Using (S6), we rewrite (S16) as

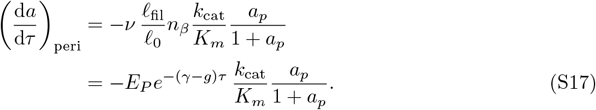

Here the number of *β*-lactamase molecules in a normal-size cell, *n*_*β*_, is multiplied with *𝓁*_fil_*/𝓁*_0_, which yields the total number of *β*-lactamase molecules in the periplasm of a filamented cell with average filament size *𝓁*_fil_. Combining this with the concentration *ν* of filamented cells, yields the expression in the last line of (S17), where *E*_*P*_ is again the concentration of *β*-lactamase in the medium if at *P* all cells had lysed simultaneously.

By our assumption of fast equilibration of the concentration gradient across the outer membrane, *a*_*p*_ on the right-hand-side of (S17) has an implicit dependence on *a* and *κ* which is given by (S8). Combining now (S15) and (S17), we obtain

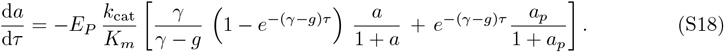

The two terms inside the rectangular brackets are the contributions due to hydrolysis in the medium and periplasm, respectively.

What remains to be done is the estimation of the CTX concentration at *τ* = 0, i.e. *t* = *t*_*P*_. This concentration is less than the initial CTX concentration, since during times up to *t*_*P*_ some CTX has already been hydrolysed in the periplasm. As we neglect lysis during the initial growth phase and further assume a constant number of enzyme molecules per unit biomass, we find that for 0 ≤ *t* ≤ *t*_*P*_, the CTX concentration of the medium changes as

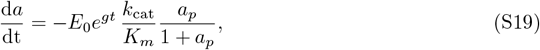

or if we are extrapolating from *t*_*P*_, so that 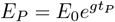,

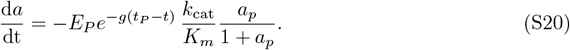

Equation (S18) and (S20) are ordinary differential equation that can be integrated numerically, given the initial CTX concentration *a*_ini_ = [CTX]_ini_*/K*_*m*_ at *t* = 0, from which the concentration at *P* can be calculated using (S20) which is then used as an initial condition for solving (S18) for *t > t*_*P*_, or equivalently, *τ >* 0. The recovery time is given by the time *τ* at which *a*(*τ*) = *a*\*, where *a*\* denotes the dimensionless CTX concentration marking the onset of OD growth. In this way, the solution of Eq. (S18) enables us to predict the recovery time, as shown in Figs. 5A and B of the main text.

### Comparing the empirical recovery times for *E. coli* K12 and REL606 with those predicted by our model

As discussed in the main text, for *E. coli* K12, the parameter *κ* is determined up to a specification of the *n*_*β*_, the number of enzyme molcules per normal-size cell, which we use as a fitting parameter. It enters the evolution equations through *E*_*P*_, the enzyme concentration at point *P*. For any given initial drug concentration, the empirically obtained OD curves furnish the average biomass growth and lysis rates, *g* and *γ*, respectively for each initial CTX concentration, as shown in Fig. S8A. As stated in the main text, we assume that recovery sets in when the external CTX concentration reaches *C* = 0.015*µ*g*/*ml. This allows us to make a theoretical prediction for the recovery time.

Figure 5A of the main text shows a comparison of the predicted recovery times *τ* with those obtained from the OD growth curves for *E. coli* K12, with *n*_*β*_ as tunable parameter. The best agreement is obtained for *n*_*β*_ = 700, which is compatible with published data. This implies that for *E. coli* K12, the parameter *κ* takes on the value *κ*_K12_ = 0.386, so that the periplasmic CTX concentration and cross-over to non-monotonic OD growth is *C*_*p*_ = 0.011*µ*g*/*ml. Note that the discrepancy of empirical results and prediction at low initial drug concentrations is due to the fact that the OD-minimum at the onset of recovery is extremely shallow and hence difficult to estimate. Moreover, note that the empirical recovery times at the onset of non-monotonic growth are consistently non-zero and of the order of a few hours across the 24 different replicates, suggesting that other physiological effects may cause an additional lag prior to recovery.

We turn next the to prediction of recovery times for *E. coli* REL606. The onset of non-monotonic growth for this strain happens at an extracellular concentration of *C* = 0.060*µ*g*/*ml. Making the natural assumption that periplasmic concentration at this point is the same as that for *E. coli* K12, i.e. *C*_*p*_ = 0.011*µ*g*/*ml, as determined above, we obtain from (S6) that *κ*_REL606_ = 5.00. We thus assume that *κ* is fixed at that value and make the additional assumption that the hydrolysis rates of CTX by AmpC are the same as for K12. The only tunable parameter is again *n*_*β*_, however due to the assumption that *κ* is fixed, an increase in *n*_*β*_, and hence enzyme concentration *E*_*p*_, must be compensated by a corresponding increase in outer membrane permeability *P*, cf. (S7).

Figure 5B shows a comparison of the predicted recovery times *τ* with those obtained from the OD growth curves for *E. coli* REL606. The best fit occurs with *n*_*β*_ = 600, which is comparable to the corresponding value obtained for *E. coli* K12. Since the number of enzymes per normal size cell are nearly equal and we have assumed that so are the hydrolization rates, the difference in the two estimated *κ*-values must be due to the difference in effective outer membrane permeability. Under the stated assumptions, our results suggest that the outer membrane permeability of *E. coli* REL606 is less than that of K12 by a factor of 15.

Figures S8C and D illustrate how the enzymatic breakdown in the two strains progresses over time. The curves shaded in red show the evolution of the extracellular CTX concentration as obtained from the numerical solution of (S18), using the empirically determined parameters for *E. coli* K12 and *E. coli* REL606. The initial concentrations chosen, along with their empirical growth, decay and lysis rates, are indicated by the circled data values in panels A and B of Fig. S8. The shading of curves in panels C and D is such that darker (lighter) shades correspond to lower (higher) initial CTX concentrations, and the concentrations plotted are normalized by the initial concentration. The curves shaded in green (purple) display the evolution of the fraction of the amount of CTX broken-down in the periplasm (medium). Note that the former curves start at non-zero values, since some CTX has already been hydrolysed during the initial growth phase. The solid curves terminate at the onset of recovery (open circles), i.e. when the external CTX concentration has been reduced sufficiently to allow for growth, and are continued by dashed lines for better visualization.

For *E. coli* K12, we see that at all times, even beyond the onset of recovery, the breakdown of extracellular CTX due to hydrolysis in the periplasm exceeds the contribution due to lysis. For times larger than the recovery time, both contributions saturate. This is in contrast to the behavior of *E. coli* REL606. Here, the contribution of lysis starts to dominate well before the onset of recovery. Further note that contrary to *E. coli* K12, the saturation of the periplasmic contribution happens well before that of the contribution due to lysis. The former is clearly due to a reduction of live biomass to a level that renders the hydrolysis in the periplasm negligible. However, by that time a sufficient amount of enzymes have been released into the medium so that hydrolysis there starts to dominate the overall hydrolyzation and saturates once all the CTX has been removed.

### Approximate solutions of the hydrolysis equations and asymptotics

In order to better understand the processes giving rise to the overall fractions of CTX in the medium vs. the periplasm, we turn to an approximate form of the hydrolysis equation (S18). Observe first that the total breakdown of external CTX due to hydrolysis in the periplasm depends on the periplasmic CTX concentration *a*_*p*_, which in turn depends on the external concentration *a* via (S6).

Depending on whether *a* ≪ *κ* + 1 or *a* ≫ *κ* + 1, the solution of (S6) become *a*_*p*_ ~ *a/*(*κ* + 1) and *a*_*p*_ ~ *a* − *κ*, respectively. We will therefore approximate the solution of (S6) as

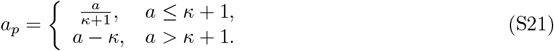

In a similar manner, we can linearize the Michaelis Menten rates as

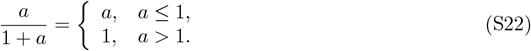

This linearization separates out the regimes where the reaction rates are proportional to the substrate concentration from those where they saturate. In terms of the drug concentration *a* in the medium, we have therefore three regions, labeled as I, II and III, as shown in Fig. S9, depending on whether *a <* 1, 1 *< a < κ* + 1 or *a > κ* + 1. The corresponding Michaelis Menten reaction rates in the medium and periplasm are shown at the bottom in blue. Regions I – III correspond to the following cases: the hydrolysis rates in the medium as well as periplasm have not saturated yet (region I), the rate in the medium has saturated but not the one in the periplams (region II) and both rates have saturated (region III). Depending on the initial and final concentration, all three regimes may be passed through during hydrolysis.

For *E. coli* K12 and REL606 the recovery concentrations are given in natural units as *a*\* = 0.206 and *a*\* = 0.823, respectively. Thus both recovery concentrations lie in region I. The circled data points in panels A and B of Fig. S8, show the initial concentration and corresponding growth and decay rates used for solving (S18) leading to the curves shown in C and D. The top horizontal axis of Fig. S8A shows the range of initial CTX concentrations in natural units *a* that *E. coli* K12 was subjected to. We can see that most of the initial concentration range lies in region I for which *a <* 1. A few data points lie in region II, i.e. 1 *< a < κ* + 1 ≈ 1.386, whose right boundary has been marked by a dashed vertical line, while a single data point lies in region III. For the REL606 strain, panel B, most of the data points lie in region II given by 1 *< a < κ* + 1 ≈ 6.

We start out by rewriting the hydrolysis equation in all three regimes as

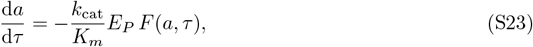

where

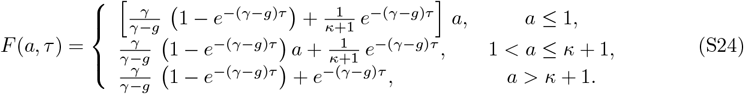

**Figure S9:**
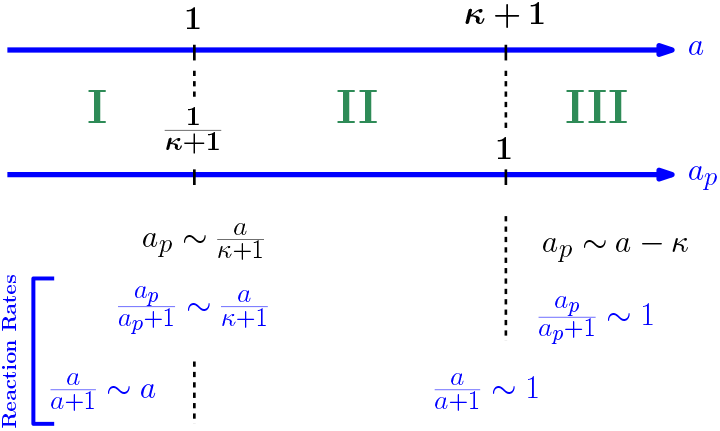
The antibiotic concentration *a* in the medium determines three dynamically distinct regions, differentiated according to whether hydrolysis in the medium as well as periplasm occurs at rates below saturation (region I), the rate in the medium has saturated (region II), or both rates have saturated (region III). Refer to text for further details.

Note that the time-dependent exponential factor *e*^−(*γ*−*g*)*τ*^ plays the role of a probability that mixes the relative rates of hydrolysis in the medium and periplasm. Specifically, consider the relative hydrolysis rate in regime I given by

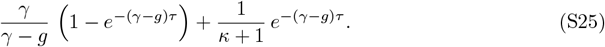

For small times (*γ* − *g*)*τ* ≪ 1, this rate is dominated by the second term 1*/*(*κ* + 1), while for large times (*γ* − *g*)*τ* ≫ 1, it is the term proportional to *γ/*(*γ* − *g*) that dominates. Further note that under conditions of intermediate decay, i.e. *γ > g*, we have that

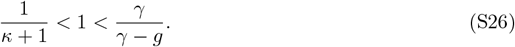

Thus the relative hydrolysis due to lysis is always larger than the corresponding periplasmic rate. Initially, most of the antibiotic is hydrolysed in the periplasm, as not sufficiently many cells have lysed yet and hence the concentration of enzymes in the medium is small. As time goes on, the relative rate of extracellular hydrolysis picks up, while that in the periplasm decreases. The cross-over time *τ*_*c*_ when the relative rates of hydrolysis in the medium and the periplams are equal is given by

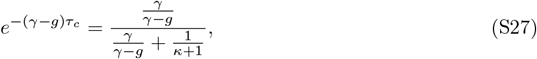

see (6). If by that time the antibiotic level has reached *a*\* or is sufficiently close to *a*\*, the hydrolysis within the medium, while becoming faster, will not have time to catch up and hence will overall contribute less to the recovery.

From (S27), we see that the cross-over time depends only on the lysis-to-decay ratio *γ/*(*γ* − *g*) and *κ*. On the other hand, the overall amount of antibiotics that has been hydrolized depends in addition on the pre-factor *k*_cat_*E*_*P*_ */K*_*m*_ in (S23), which can vary independently of *τ*_*c*_. For example, if *E*_*P*_ is large, then by the cross-over time *τ*_*c*_ most of the antibiotic will have been hydrolised in the periplasm and hence the contribution via lysis will be less, while if *E*_*P*_ is sufficiently small, hydrolysis due to lysis will dominate the process leading to recovery.

We now turn to a solution of (S23) in order to determine the contributions of the hydrolysis in the medium and periplasm to the overall reduction of extracellular CTX. For simplicity we will consider only solutions in region I, i.e.

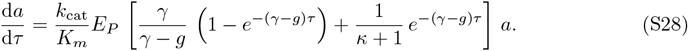

Note that in all three regions the equations are linear and hence can be solved exactly. The analysis where the initial concentration starts out in one of the other two regions will thus be similar.

For region I, it is convenient to introduce the following dimensionless quantities

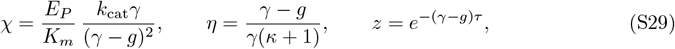

where *χ* is a ratio of the rates of hydrolysis and effective biomass decay, *η* is the ratio of relative hydrolysis rates due to activity in the periplasm and medium, describing the relative privatization of CTX hydrolysis, while *z* is a reparametrization of the time, with *z* = 1 corresponding to *τ* = 0 and *z* decaying to zero as *τ*. The fraction of initial CTX that remains at time *τ* → ∞ is then given by

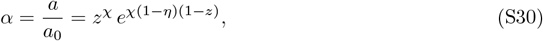

where the dependence on *τ* is via *z* as given in Eq. (S29). Alternatively, it is useful to think of Eq. (S30) as defining *z* in terms of *α*.

Note that *a*_0_ − *a* is the amount of CTX hydrolysed at time *τ*. Integrating (S28) and dividing through by *a*_0_, we find

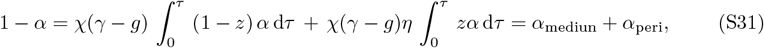

where the first and second terms containing integrals are the fractions of the CTX hydrolysed in the medium and periplasm at time *τ*. We denote these fractions as *α*_medium_ and *α*_peri_, respectively. It then follows that

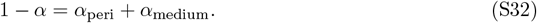

Equation (S32) provides the fractional contributions of medium and periplasm to the total amount CTX broken-down so far. Substituting the explicit form of *α* given by (S30), we can determine *α*_peri_ as

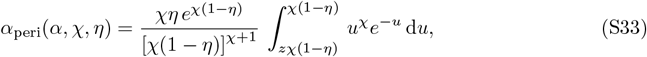

where the dependence on *α* is via *z* according to (S30). The corresponding expression for *α*_medium_ follows from (S32) and is given as *α*_medium_(*α, χ, η*) = 1 − *α* − *α*_peri_(*α, χ, η*). Equation (S33) is our central result. It shows how the fraction of CTX broken down due to hydrolysis in the medium depends on time via *α*, and on the parameters *χ* and *η* describing the relative hydrolysis and privatization rates, respectively.

The case where equal amounts of CTX is hydrolized in the medium and periplasm, corresponds to a choice of parameters *α, χ* and *η* such that

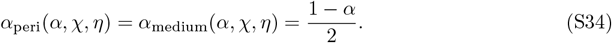

Equations (S33) together with (S34) define a surface in this space of parameters, which divides it into regions where either hydrolysis in the periplasm dominates over that in the medium or vice versa.

### Recovery time in the limit of fast *β*-lactam degradation

A simple explicit expression for the recovery time *τ*_rec_ defined by (10) can be obtained in the limiting case when Γ is sufficiently large, so that recovery happens very quickly, and therefore the exponential terms in equation (3) can be set to unity. In this limit, the drug concentration declines exponentially at rate Γ*/*(*κ* + 1) and the recovery time is given by

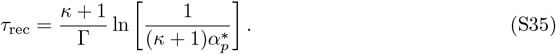

The recovery time vanishes when the initial concentration equals 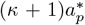. With decreasing privatization, *τ*_rec_ increases, reaching a maximum at 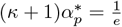 and subsequently approaching a limiting value of 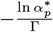 at *κ* = 0 (see Fig. 6B).

